# Hierarchical Bayesian models of transcriptional and translational regulation processes with delays

**DOI:** 10.1101/2021.08.16.456485

**Authors:** Mark Jayson Cortez, Hyukpyo Hong, Boseung Choi, Jae Kyoung Kim, Krešimir Josić

## Abstract

**Motivation:** Simultaneous recordings of gene network dynamics across large populations have revealed that cell characteristics vary considerably even in clonal lines. Inferring the variability of parameters that determine gene dynamics is key to understanding cellular behavior. However, this is complicated by the fact that the outcomes and effects of many reactions are not observable directly. Unobserved reactions can be replaced with time delays to reduce model dimensionality and simplify inference. However, the resulting models are non-Markovian, and require the development of new inference techniques.

**Results:** We propose a non-Markovian, hierarchical Bayesian inference framework for quantifying the variability of cellular processes within and across cells in a population. We illustrate our approach using a delayed birth-death process. In general, a distributed delay model, rather than a popular fixed delay model, is needed for inference, even if only mean reaction delays are of interest. Using *in silico* and experimental data we show that the proposed hierarchical framework is robust and leads to improved estimates compared to its non-hierarchical counterpart. We apply our method to data obtained using time-lapse microscopy and infer the parameters that describe the dynamics of protein production at the single cell and population level. The mean delays in protein production are larger than previously reported, have a coefficient of variation of around 0.2 across the population, and are not strongly correlated with protein production or growth rates.

**Availability:** Accompanying code in Python is available at https://github.com/mvcortez/Bayesian-Inference.

**Contact:** @gmail.com

## 1 Introduction

Cellular processes are inherently variable, both in time, and across individuals in a population. Characterizing such variability, and quantifying how features of cellular processes covary with phenotype and genotype is essential to understanding cellular behaviors. The estimation of such covariability often requires the analysis of time series of measurements from different cells across a clonal population [Hasenauer *et al*., 2010; Zechner *et al*., 2014; Koeppl *et al*., 2012]. Hierarchical modelling provides a systematic framework to analyze such population level data, and characterize both cell-to-cell and within cell variability [Heydari *et al*., 2016; Tonner *et al*., 2020]. This framework improves the robustness of the estimates of parameters that describe processes within individual cells by assuming that these parameters follow an underlying population distribution [Gelman *et al*., 2013; Congdon, 2020].

While elegant, hierarchical models are often high dimensional and computationally intractable [Zechner *et al*., 2014]. It is therefore necessary to base such models on compact, yet interpretable descriptions of single cell processes. One way to obtain such descriptions is through coarse–grained models of cellular dynamics: Many cellular processes are composed of a sequence of substeps, which are often not of interest in themselves. In models, we can often replace such reaction chains by a single reaction, at the expense of introducing a delay [Barrio *et al*., 2013; Leier *et al*., 2014; Gomez *et al*., 2016]. For example, the production of a regulator protein can be described as a sequence of reactions including transcription, translation, and post-translational steps [Golding *et al*., 2005; Kaern *et al*., 2005]. This entire process can also be described more coarsely as a single reaction, which, once initiated, takes a random time to complete [MacAdams *et al*., 1995].

The introduction of delays results in a non-Markovian process, making inference challenging. Stochastic delay differential equations are often used to model processes with delays. The exact stationary probability densities of these equations have been used to identify the components of the systems under study [Frank *et al*., 2001, 2003]. Progress has also been made by using artificial neural networks to approximate delay chemical master equations [Jiang *et al*., 2021], an approach which works well for the inference of parameters of a birth-death process with a delayed death reaction. Likelihood-based inference using the chemical Langevin equation descriptions of the delayed process [Heron *et al*., 2007], and linear noise approximations [Calderazzo *et al*., 2019] have also been used. These approaches, however, are effective only when molecule counts are high and stochastic differential equations accurately capture system dynamics [Gupta *et al*., 2014]. Choi et al. [2020] developed an alternative Bayesian approach using non-Markovian models to develop inference algorithms for rate and delay parameters in common biochemical reactions. This approach works well with synthetic and experimental data, and is effective even when molecule counts are low because the model is based on the chemical master equation. However, it relies on treating measurements from different cells as independent, identically distributed observations of a single cellular process, thus compounding uncertainty in parameter estimates with variability across the population of cells.

To address this problem, we develop a hierarchical, non-Markovian biochemical reaction model, and derive associated likelihood functions to characterize variability within and across cells. This allows us to develop a sampling algorithm to simultaneously estimate the posterior distribution of parameters characterizing processes within individual cells, as well as the distribution of these parameters across the population. We demonstrate the advantages and shortcomings of our approach using a delayed birth-death process, which, although it may not fully describe the underlying biophysical processes, captures the main effects of protein production, and can serve as a building block for more complex systems. When birth delays vary between individual reactions, using a popular fixed birth delay model for inference leads to a biased estimate of mean reaction delays. Hence, to accurately describe reaction timing, we next develop an inference method based on a distributed delay model, which allows for the estimation of model parameters for individual cells in the population, and the analysis of correlations between different model parameters. We use our approach to infer the production rate and delay in the synthesis of yellow fluorescent protein (YFP) from time-lapse microscopy measurements. Across the population estimated production rates are highly variable with a coefficient of variation (CV) of around 0.5, while mean delay times have a CV of around 0.2. Unexpectedly, we find that delay is not strongly correlated with production rate or growth rate. Our mean delay estimates are higher than those reported previously [Choi *et al*., 2020]. Using synthetic data we show that this difference can be explained by a bias in mean delay estimates when assuming all cells are identical, and illustrate that the parameters inferred using a hierarchical approach reproduce the experimentally observed YFP dynamics in individual cells. We also show that hierarchical inference performs better than a non-hierarchical analog in estimating delay distribution variance, especially when data is coarsely-sampled. Our non-Markovian, hierarchical Bayesian inference framework thus provides an effective tool for identifying cell-to-cell and within cell variability. While our examples are limited to a birth-death process, our approach is general, and can be applied to more complex biochemical processes.

## 2 Methods

### 2.1 Derivation of the likelihood function given multi-cell observations

Consider a chemical reaction network which describes the evolution of *u* species, *Y*_1_, *Y*_2_, …, *Y*_*u*_, through a set of *v* chemical reactions, *R*_1_, *R*_2_, …, *R*_*v*_. We represent this system as

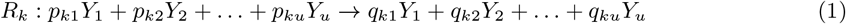

where *p*_*kj*_ and *q*_*kj*_ are stoichiometric constants. For each reaction *R*_*k*_, there is a stochastic rate constant, *θ*_*k*_, and function, *h*_*k*_((*y*(*t*), *θ*_*k*_), that describes the instantaneous hazard of reaction *R*_*k*_ occurring under some kinetic law, where *y* (*t*) = (*y*_1_ (*t*), *y*_2_ (*t*), …, *y*_*u*_ (*t*)) is the molecular count of all chemical species at time *t*.

We assume that the reactions in Eq. (1) are not completed instantaneously. We denote by *t*_initial_ and *t*_final_ the initiation and completion times of a reaction, respectively, so the time to completion, or delay, is given by *t*_final_ − *t*_initial_, and it can be either fixed or a random variable. With each reaction, *R*_*k*_, we can thus associate a delay measure *η*_*k*_, with support on [0, ∞). We assume that the delay distribution for reaction *k* does not depend on time or the state of the system, and only depends on a set of *l*_*k*_ parameters 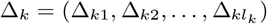.

Schlicht et al. [2008] have proven the existence of reaction completion propensities defined by

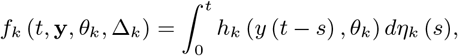

which serves as the effective rate of reaction at a given time *t*. Integrating with time, we have 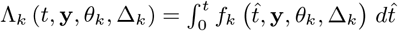, and 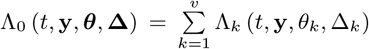, where **y** denotes the trajectory of all chemical species from time 0 to time *T*, and ***θ*** and **Δ** denote the vector of all the rate and delay parameters, respectively. Building on this result, Choi et al. [2020] derived likelihood functions for reaction rate and delay parameters for an individual stochastic process described by Eq. (1).

Suppose that the process **y** is fully-observed over the time interval [0, *T*], let *r*_*ki*_ be the number of reactions of type *k* that are completed in the time subinterval (*i, i*+1], and let 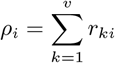. Every reaction occurring within the subinterval (*i, i* + 1] is associated to reaction time and type (*t*_*ij*_, *k*_*ij*_), *j* = 1, 2, …, *ρ*_*i*_. The likelihood function for the parameters ***θ*** and **Δ** is given by

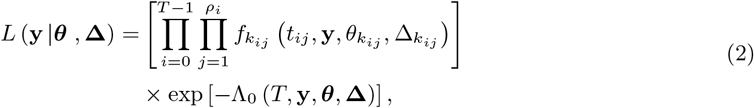

which is analogous to the likelihood provided by Boys et al. [2008] for a system without reaction completion delays.

Suppose that the process **y** is observed at discrete times *t* ∈ {0, 1, …, *T* − 1, *T*}, so that only the observations **y**_**d**_ = (*y* (0), *y* (1), …, *y* (*T* − 1), *y* (*T*)) are available. Then the completion propensity, *f*_*k*_, can be approximated by its average, 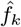, on each unit time interval, (*i, i* + 1], obtained from linearly interpolating the reaction hazard, *h*_*k*_, between observations [Boys *et al*., 2008],

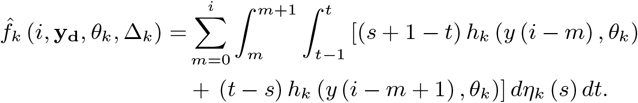

Conditioned on the entire history of the system up to time *i*, the number of reactions of type *k* that completed within (*i, i*+1] are independent and follows a Poisson distribution with mean 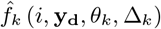. Hence, the likelihood given by Eq. (2) can thus be approximated by

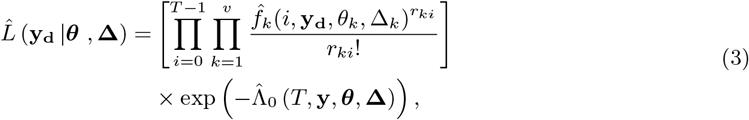

where 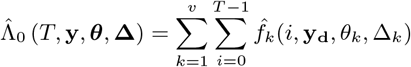 [Gupta *et al*., 2014].

We assume that we are observing a population of *N* cells, leading to distinct sequences of observations (**y**_**1**_, **y**_**2**_, …, **y**_**N**_) on the time interval [0, *T*] (one for each cell). We observe the outcomes of the same set of reactions in each cell, but the parameters characterizing each reaction can differ between cells. Each reaction, *R*_*k*_, is endowed with a rate constant *θ*_*nk*_, so that reactions in cell *n* are characterized by the rate vector *θ*_*n*_ = (*θ*_*n*1_, *θ*_*n*2_, …, *θ*_*nk*_, …, *θ*_*nv*_). If a reaction *R*_*k*_ in cell *n* is delayed, then we also associate to that reaction the delay parameters 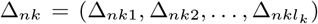, and denote the set of all delay parameters for cell *n* by Δ_*n*_ = {Δ_*nk*_}. We denote by ***θ*** the collection {*θ*_*n*_} of all rate constants and by **Δ** the collection {Δ_*n*_} of parameters that define all delay measures. In the Supplementary Material, we provide a detailed description in the cases that delays are described by a Gamma or a Dirac delta distribution.

Suppose that the process **y**_**n**_ is fully observed over the interval [0, *T*]. We can then define the total likelihood

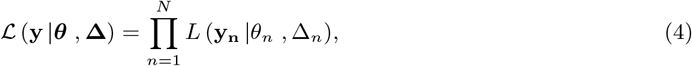

which is the product of likelihoods given in Eq. (2) for all *N* individuals. If only the discrete-time observations **y**_**d**,**n**_ = (*y*_*n*_ (0), *y*_*n*_ (1), …, *y*_*n*_ (*T* − 1), *y*_*n*_ (*T*)) are available, we can approximate the total likelihood by

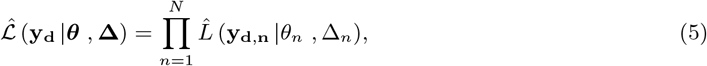

which is a product of likelihoods given in Eq. (3). As before, the computation of this approximate likelihood, 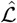, requires information on the number of completed reaction of type *k* on each interval (*i, i* + 1] for each cell *n*, which we write as *r*_*nki*_.

### 2.2 A hierarchical Bayesian model of a cell population

Bayes’ Theorem and the independence of observations enable the factorization of the joint posterior of parameters and hyperparameters, and hence allow us to take a multilevel approach (i.e., a hierarchical modeling approach) to inference. Just as the observations **y**_**d**,*n*_ depend on the rate and delay parameters, we can assume that the individual-level parameters *θ*_*nk*_ and Δ_*nkl*_ follow underlying distributions which are themselves characterized by hyperparameters, 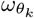 and 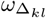, respectively. We also assume that these individual parameters are independent for given hyperparameters, as we are observing cells that are not closely related.

Denoting the collection 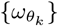 and 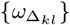 of rate and delay hyperparameters respectively as *ω*_*θ*_ and *ω*Δ, the approximate likelihood expression given in Eq. (5), and Bayes’ Theorem allow us to write the posterior over the parameters characterizing the biochemical reaction network, to reflect the sequence of observation-parameter and parameter-hyperparameter dependencies as

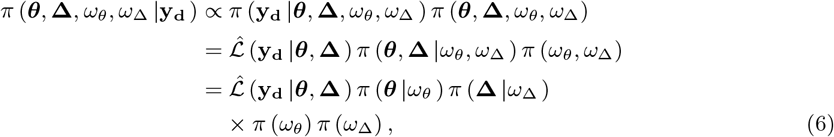

with 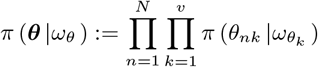 and

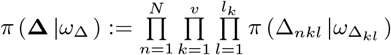 serving as priors for individual rate and delay parameters, and *π* (*ω*_*θ*_) and *π* (*ω*_Δ_) as the hyperpriors.

### 2.3 Inference with a hierarchical model for heterogeneous cell populations

We next describe an MCMC algorithm to generate samples from the posterior distribution of the model parameters (***θ*, Δ**) and corresponding hyperparameters (*ω*_*θ*_, *ω*_Δ_). The priors and hyperpriors capture our previous knowledge about the variability of the parameters across the population. As rate parameters are positive, we use gamma distributions as priors, 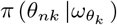. Thus for every reaction *k*, the set of hyperparameters for the corresponding reaction rate is 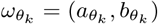, where 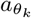 and 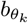 are the shape and rate parameters respectively of a gamma distribution. If the reaction propensity is separable, as in the case of mass-action kinetics where the hazard function can be factored as *h*_*k*_ (*y*_*n*_ (*t*), *θ*_*nk*_) = *θ*_*nk*_*g*_*k*_ (*y*_*n*_ (*t*)), the gamma distribution defines a conjugate prior for the parameters *θ*_*nk*_ [Wilkinson, 2011].

As is typical of hierarchical sampling approaches, our algorithm iteratively produces samples of individual parameters and integrates the result across an ensemble of cells to produce a sample of the hyperparameters that characterize the population distribution. The updated population distribution is then used to generate new samples of individual cell parameters, and the process repeats. To sample from the posterior distribution given by Eq. (6), we use Gibbs sampling: For every individual cell, *n*, we obtain samples for *θ*_*n*_ and Δ_*n*_ from their conditional posterior distributions by using the Metropolis-Hastings algorithm [Hastings, 1970; Metropolis *et al*., 1953]. As described by Choi et al. [2020], knowledge about the number of completed reactions, *r*_*nki*_, is needed in the sampling of these individual parameters. Since the discrete-time measurements are not sufficient to uniquely determine the number of reactions, we sample *r*_*nki*_ in each Gibbs step. To do so, we follow the block-updating strategy described by Boys et al. [2008], and infer the number of reactions in each interval (*i, i* + 1] through the Metropolis-Hastings algorithm with a random-walk chain. In this scheme, a proposal is generated by augmenting the current value by a random variable from the Skellam distribution [Boys *et al*., 2008; Johnson *et al*., 1969]. Once samples of both rate and delay parameters for all *N* cells are obtained, we sample the hyperparameters *ω*_*θ*_ and *ω*_Δ_ using the Metropolis-Hastings algorithm and the individual-level parameters as data in the population-level sampling.

We give a full description of the MCMC algorithm in the Supplementary Methods. There we also derive the likelihoods and resulting posterior distributions for a stochastic birth-death process with birth delays that we use as the main example in the following.

## 3 Results

### 3.1 Heterogeneous fixed delays can be accurately estimated

We first demonstrate the inference process using synthetic data generated from a population of stochastic birth-death processes with fixed birth delays [Heron *et al*., 2007; Bel *et al*., 2009; Gupta *et al*., 2014; Calderazzo *et al*., 2019],

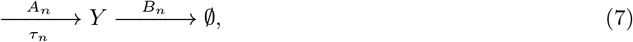

where *n* = 1, 2, …, *N* and *N* is the number of cells (see Eq. S14 and S15 for the likelihood function and full posterior, respectively).

Although simple, this delayed birth-death process can be used to model the dynamics of chemical species, such as proteins, that are produced through a sequence of reactions. For simplicity, we refer to the product, *Y*, in Eq. (7) as a protein. We assume that protein expression is Poissonian rather than bursty [Thattai and van Oudenaarden, 2001; Munsky *et al*., 2012]. We expect this assumption will hold with high gene copy numbers, as in the experimental systems we will study subsequently. We first consider fixed birth delays, *τ*_*n*_, which are constant across reactions within a cell, but may vary between cells. Later we consider distributed delays that vary between reactions within and between cells. In the generative model (Fig. **S1**a), we assume that, across the population, production rates, *A*_*n*_, degradation rates, *B*_*n*_, and the fixed birth delays, *τ*_*n*_, follow a gamma distribution. We also assume that protein numbers, *Y*, are exactly measurable at discrete times. The population is induced at time *t* = 0. Assuming the cognate promoters are not leaky, the production rate in each cell, *n*, then changes from 0 to the fully induced values, *A*_*n*_, instantaneously at time of induction. Decrease in protein count is due to growth-induced dilution or enzymatic degradation, and is described by an instantaneous death process with rate *B*_*n*_.

Each cell in the population is thus described by three parameters: the rate constants *A*_*n*_, *B*_*n*_, and the fixed birth delay time *τ*_*n*_. To generate synthetic population data, we sampled 40 triplets of these parameters from their corresponding gamma distributions and used them to generate 40 realizations of the birth-death process using the delayed Gillespie algorithm [Barrio *et al*., 2006]. To mimic experimental data, we subsampled the resulting trajectories by recording the molecular counts at unit time intervals (Fig. **S1**b).

We next compared the parameters estimated using our hierarchical inference algorithm with those used to generate the synthetic data. We first used non-informative, rational hyperpriors for all hyperparameters. When full trajectories (Fig. **S1**b) are used, *A*_*n*_, *B*_*n*_, and *τ*_*n*_ were accurately estimated (Fig. **S1**c and d; blue dots). On the other hand, for the initial segment of the trajectories (red box in Fig. **S1**b), the protein count, *Y*_*n*_(*t*), was small and thus few deaths (*B*_*n*_ · *Y*_*n*_(*t*)) occurred. Hence death rates were underestimated (Fig. **S1**c; orange dots). Birth rates were also underestimated to compensate for the low death rates (Fig. **S1**c; orange dots), but delays were estimated well (Fig. **S1**d; orange dots). If we assumed that the death rates, *B*_*n*_, are known, both *A*_*n*_ and *τ*_*n*_ were accurately estimated even with the shorter trajectories (Fig. **S1**e). Since dilution rates can be measured by tracking cell growth, death rates can be estimated from experimental data [Choi *et al*., 2020]. Hence, inferring delays and birth rates from realistic amounts of data (e.g., 40 min in this case) is possible [Cheng *et al*., 2017] if the *B*_*n*_ can be directly measured.

**Fig. S1:**
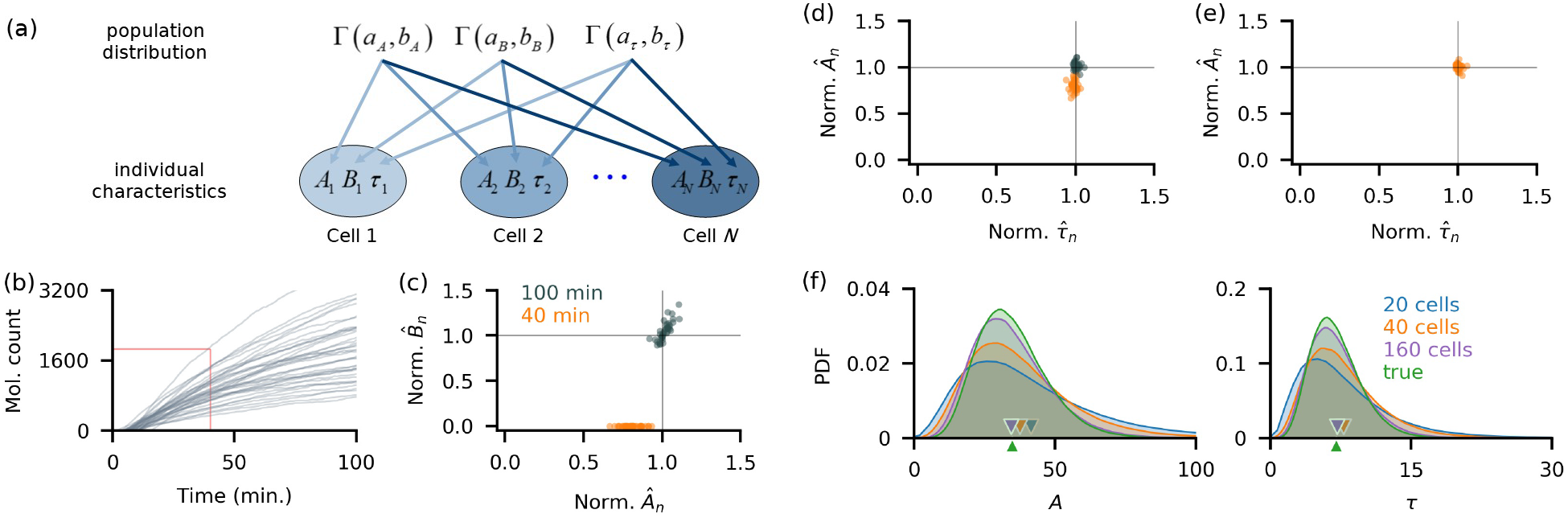
Fixed delays and production rates are accurately estimated both at the population and individual levels. (a) Individual birth-death processes, are described by the production rate, *A*_*n*_, the degradation rate, *B*_*n*_, and the fixed birth delay time, *τ*_*n*_. We assumed these parameters follow Gamma distributions. (b) Simulated trajectories of birth-death processes with fixed birth delays subsampled every minute. Individual parameters were sampled from the following distributions similar to those estimated from YFP synthesis trajectories in a previous study [Choi *et al*., 2020]: *A*_*n*_ ∼ Γ(8, 0.23), *B*_*n*_ ∼ Γ(9, 625), and *τ*_*n*_ ∼ Γ(7, 1). To estimate all three parameters per cell, we implemented the algorithm initially using the first 40 min (red box in (b)), and 100 min of observation. In panels c-f we divided each estimate with the true parameter value, so that a perfect match corresponds to 1. (c-d) With 100 min of data, all rates (c) and delays (d) are accurately estimated (blue dots). However, 40 min of data lead to underestimates of the death rates, *B*_*n*_ (c). The birth rates, *A*_*n*_, were underestimated to compensate for the low death rate estimates (c). Estimates of the fixed delay times were still accurate (d). (e-f) When the degradation rates, *B*_*n*_, were assumed known, the posterior means for each cell were close to the true parameter values (e). Population-level posterior densities of both the growth rate, *A*, and delay time, *τ*, were wider than the true densities, but their means (triangular markers) were close to the true value (f). The inferred population distributions improved with an increase in the number (from 20 to 160) of observed realizations of the stochastic processes.

Estimates of the distributions of the parameters across the population improved with the number of observed trajectories (Fig. **S1**f), showing that the algorithm is consistent, characterized by the convergence of posteriors to the true distributions with more data. This is further evidenced by the improvement in accuracy and precision of the hyperparameter estimates with an increase in the number of observed trajectories (See Fig. S1). We therefore concluded that our hierarchical algorithm can be used to simultaneously infer reaction parameters for individual cells, as well as the distribution of these parameters across a population from realistic amounts of data [Cheng *et al*., 2017].

Parameter identifiability analysis is well-established for deterministic, but not stochastic systems [Browning *et al*., 2020]. *Practical identifiability* – the question of whether a parameter can be estimated from a finite amount of noisy data with accuracy beyond that given by the prior distribution – is most relevant in practice [Raue *et al*., 2009; Hines *et al*., 2014]. Comparing the prior and posterior distributions in our examples with synthetic data shows that the parameters of birth-death processes with delays are practically identifiable, at least in the parameter ranges we considered. However, even in the limit of infinite sampling rate the posterior distributions over the individual parameters will not converge to point masses at the true values when the time of observations is finite. This is because the number of observed reactions over a finite interval is finite, regardless of sampling frequency. Whether the individual parameter estimates converge to their true values as the observation window diverges, or whether the hyperparameter estimates converge to their true values as the number of cells diverges, as suggested by Fig. **S1**f and S1, are open questions.

### 3.2 A distributed delay model provides accurate estimates of mean delay times

In biological systems, delay is usually distributed rather than fixed. However, since we are sometimes interested only in mean reaction delays, we asked whether we can use a simple, hierarchical fixed delay model to estimate reaction parameters even in situations when delays are distributed. To answer this question we generated trajectories using a model in which individual reaction delays followed a gamma distribution, *τ*_*n*_ ∼ Γ(*α*_*n*_, *β*_*n*_) (Fig. **S2**a), with parameters, *α*_*n*_ and *β*_*n*_, that could differ between individual cells in the population (Fig. **S2**b). We chose three sets of parameters *α*_*n*_ and *β*_*n*_ so that for each set the mean delay across the population was the same (See Table S1), while the variances of the *individual* delay distributions differed between the sets: 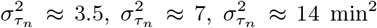. In each case, we simulated 40 trajectories, each with 40 min of observations at 1 min intervals (Fig. **S2**c).

When we applied our inference algorithm using a hierarchical fixed delay model to this data, we observed that both mean delay times and birth rates were underestimated, and that this bias increased with the variance of the individual delay distributions (i.e., farther from the fixed delay) (Fig. **S2**d-f; orange dots). This is consistent with the findings of Josić et al. [2011] who showed that distributed delays can accelerate signaling in genetic networks by reducing the time for a process to reach threshold compared to systems with fixed delay. In the present case, since delay times were distributed, the earliest detectable signal after induction was likely to be observed before the mean delay time. A model with fixed delay will interpret the first observation of the product, *Y*, as the delay time to the completion of the first birth reaction after initiation. As a consequence, this will lead to an underestimate of each delay. This implies that previous inference results obtained with fixed delay models need to be interpreted carefully [Gopalakrishnan *et al*., 2011; Wang *et al*., 2012; Mehrkanoon *et al*., 2014], as even the inference of mean delay times generally requires an algorithm based on a distributed delay model.

We next asked whether parameters of a population of birth-death processes with distributed delays can be accurately estimated using a distributed delay model (Fig. **S2**b). Using the same set of non-informative rational hyperpriors as in the fixed delay model resulted in accurate estimates of the individual production rates, *A*_*n*_, and mean delay times, 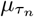 (Fig. **S2**d-f; blue dots). The delay estimates were further improved by using a non-informative maximal data information prior (MDIP) [Pradhan *et al*., 2011; Zellner, 1991] over the delay hyperparameters (Fig. **S2**d-f; red dots). The MDIP incorporates the dependence structure of the parameters (*α*_*n*_, *β*_*n*_) of the Gamma delay distribution [Moala *et al*., 2013]. In addition, the MDIP emphasizes the information supplied by the data density rather than that of the prior, thereby providing weaker influence than the information given by the data itself. At the population level, the distributions of both the production rate, *A*, (Fig. **S2**g-i) and delay time, *τ*, (Fig. **S2**j-l) were similar for both non-informative hyperprior choices and closely matched the true distributions. Henceforth, we used the MDIP as the default non-informative delay hyperprior. Using hyperpriors that capture pre-existing knowledge about the delay distribution also improves estimates of reaction rates and delay parameters (See Supplementary Methods and Fig. S2).

**Fig. S2:**
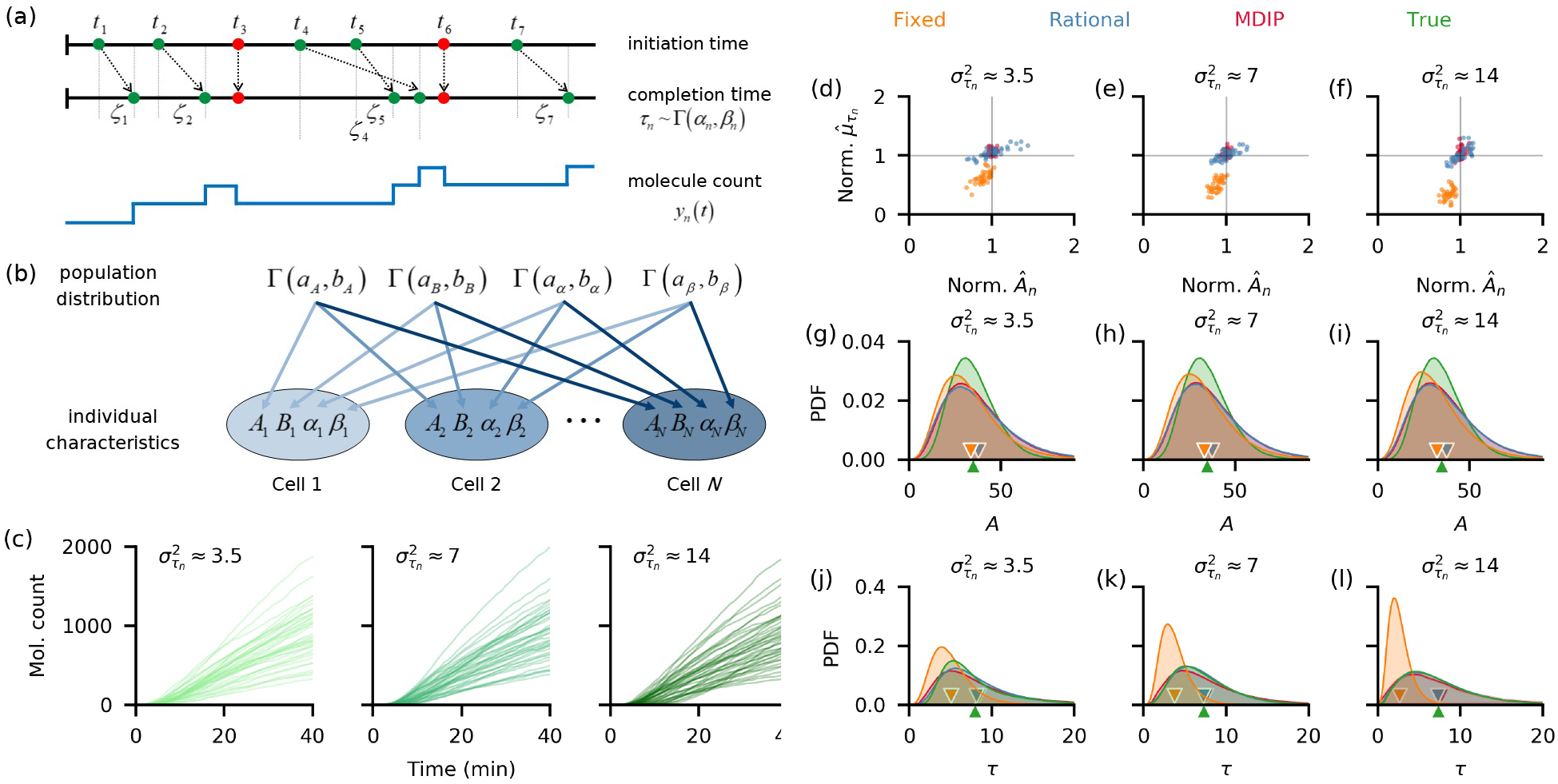
A hierarchical distributed delay model leads to accurate estimates of distributed delays, while a fixed delay model underestimates delays. (a) A birth reaction in each cell *n* is initiated at time *t*_*i*_ and completed after a delay, *ζ*_*i*_. Each delay is a realization of the random variable *τ*_*n*_ which follows a Gamma distribution with parameters (*α*_*n*_, *β*_*n*_). Death reactions (red) are instantaneous. (b) The generative model for a birth-death process with distributed delays. Each individual process, *n*, is described by four parameters: the production rate, *A*_*n*_, the degradation rate, *B*_*n*_, and the two parameters describing the delay distribution, (*α*_*n*_, *β*_*n*_). All parameters follow Gamma distributions, with respective hyperparameters. (c) To generate trajectories, we fixed a set of production and degradation rates, *A*_*n*_, and *B*_*n*_, and chose three different sets of delay parameters *α*_*n*_ and *β*_*n*_. Mean delays were equal for all cases while delay variances within a cell were approximately equal across each population, but differed between the three cases (See Table S1.). For each parameter, set we simulated 40 trajectories that were subsampled every minute. (d-f) Across all data sets, both the production rates *A*_*n*_ and mean delay times, 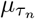, were underestimated when we used the fixed delay model (orange dots), but were accurately estimated with the distributed delay model with either rational hyperpriors (blue dots) or MDIP (red dots) over the delay hyperparameters. With the fixed delay model, the bias in the mean delay estimate increased with within-cell delay variance, 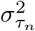. For comparison we, normalized the estimated parameters by dividing with the true values. Here, we assumed *B*_*n*_ is known. (g-i) The estimated population distributions of the production rates were similar for the distributed delay models, with means (triangular markers) close to those of the true distributions. The posterior obtained with the fixed delay model gave a slight underestimate of the mean population production rate. (j-l) The pooled posterior delay distributions obtained using the distributed delay model matched the true distribution. The bias in the pooled posterior delay distributions obtained using the fixed delay model increased, and the estimated mean population delay (orange triangular marker) approached zero, as delay variance, 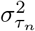, increased.

**Fig. S3:**
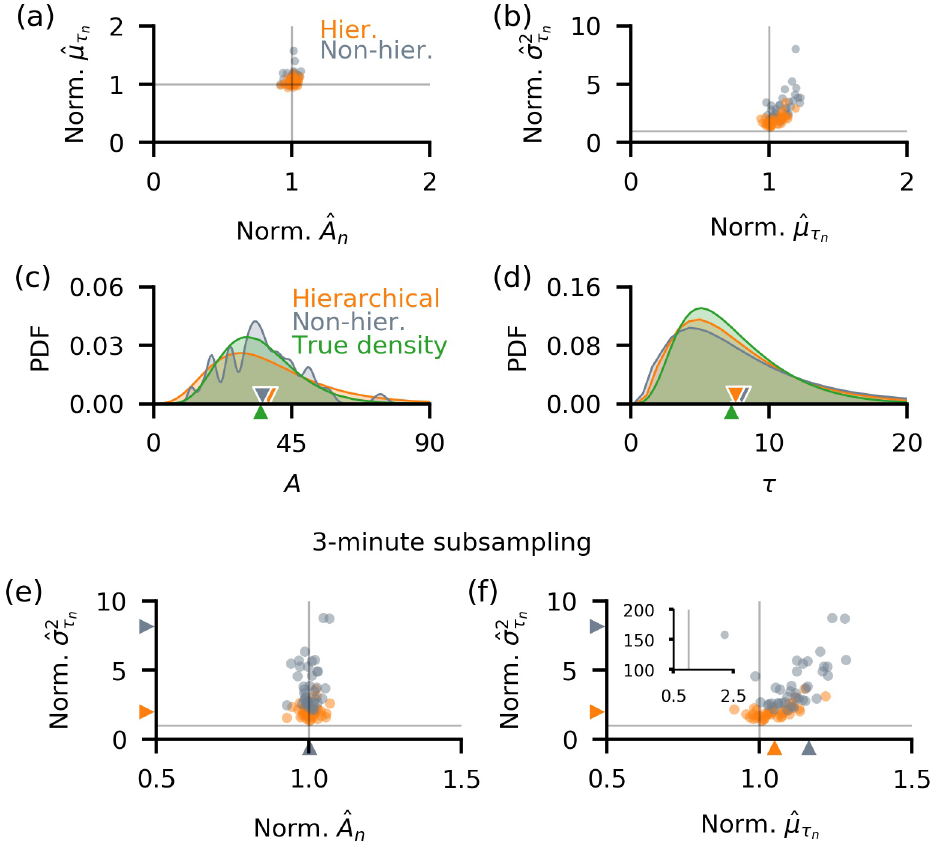
Hierarchical inference outperforms non-hierarchical inference and leads to better estimates of delay variances. Panels (a-b) show the individual parameter estimates normalized by the true values. (a) Although individual production rate estimates were similar for both approaches, the hierarchical model produced better estimates of mean delays with fewer outliers. (b) Delay variances are similarly better estimated by the hierarchical model. (c-d) Comparison of inferred population-level distributions of production rates, *A*, and delay times, *τ*, exhibit the same advantages of the hierarchical model. (e-f) In model implementation using 3-minute subsampled trajectories, the accuracy of inferred production rates, *Â*_*n*_ (e), and mean delay times, 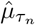 (f), is similar, but the hierarchical model has a smaller bias, and produces fewer outlying estimates. With non-hierarchical inference, there was an extreme outlier which corresponded to overestimates of 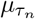 and 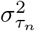 (f inset). The hierarchical model provided better estimates of the individual delay variances. We used non-informative priors over parameters in all cases (See Supplementary Methods).

Using a fixed delay model to infer mean delay times introduces bias in inference when delay, in itself, varies. We conclude that a distributed delay model should be used for inference whether only the average delay, or more detailed information about the delay distribution, like higher-order statistics [Blyuss and Kyrychko, 2010; Kyrychko *et al*., 2013], are of interest. An algorithm based on the distributed delay model provided accurate estimates of mean delays whether individual delay distributions were wide (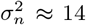 in Fig. **S2**c) or point-masses (see Fig. S3). Although the population level posteriors (Fig. **S2**j-l) were slightly wider than the true densities, the mean delay times were accurately estimated. Thus, our hierarchical model is robust to changes in hyperparameters, and is applicable to populations with varied characteristics.

### 3.3 Hierarchical inference outperforms non-hierarchical inference

Hierarchical methods typically shrink individual parameter estimates leading to more robust estimates of population-level parameters, but also increase model complexity [Congdon, 2020]. As an alternative, we can infer reaction parameters individually from each observed trajectory and estimate the population-level parameters from the collected individual estimates. This approach is typically less robust than hierarchical inference, but is also less computationally expensive. We therefore asked what advantages hierarchical inference offers over a non-hierarchical approach with the types of data that motivated this study.

To compare hierarchical and non-hierarchical inference, we again considered measurements from a collection of birth-death processes with distributed birth delays (1-min subsampled trajectories with 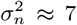 in Fig. **S2**c), and used non-informative priors for both cases (See Supplementary Methods). While the estimates of the individual birth parameters, *A*_*n*_, were similar for both models, the hierarchical model provided better estimates of the mean delay times, 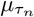 (Fig. **S3**a). Although individual delay variances 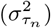 were overestimated in both cases, the hierarchical model provided a better estimate (Fig. **S3**b). The hierarchical model also gave better estimates of the production rates and delay times (Fig. **S3**c and d) over a range of single cell and population level parameters we used to generate the data (See Fig. S4).

The overestimates of 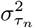 were due to the strong correlation between the inferred parameters *α*_*n*_ and *β*_*n*_, (See Fig. S5d-f),which consequently made an accurate estimation of both individual delay mean and variance difficult. A similar observation has been made by Choi et al. [2020] for estimates from single trajectories. They proposed pooling multiple recordings to increase estimate accuracy, implicitly assuming that all cells in the population are identical. The resulting estimates captured the mean of the parameter across the population. However, this approach may lead to biased parameter estimates when cell populations are heterogeneous, as we show below (See also Fig. S6). The hierarchical model, which performs such pooling without assuming that cells are identical, returns estimates that better capture the parameter central tendencies (See Fig. S6).

When generating synthetic data we used sampling rates consistent with those obtainable using timelapse fluorescence microscopy [Golding *et al*., 2005; Garcia *et al*., 2013; Chen *et al*., 2015]. In this range, the frequency of measurements has a strong impact on the accuracy of individual parameter estimates. We thus expected that the non-hierarchical model performs worse than its hierarchical counterpart when sampling frequency is low, but both produce similar individual cell estimates at high sampling frequencies. We tested this prediction by generating data from 20 individual 60 min trajectories, each sampled once per 3 min (in contrast to 1 sample per minute we used previously). The hierarchical and non-hierarchical models produced similar production rate estimates (Fig. **S3**e), but the mean delay times (Fig. **S3**f) and individual delay time variances (Fig. **S3**e and f) were estimated more accurately with the hierarchical model. We further explore this in the Supplementary Material where we show that the hierarchical model yielded consistent results at a range of sampling frequencies: individual parameter estimates were accurate, delay variance estimates did not diverge even with low-resolution data (see Fig. S7a-e), and KL-divergence between the the delay population posterior and the true density remained small, and depended weakly on sampling frequency (see Fig. S7f)). The non-hierarchical model, on the other hand, produced estimates which decreased in accuracy with the increase in sampling interval, produced delay population posteriors that were far from the true distributions, as well as extreme outlying individual parameter estimates (see Fig. S7a-e).

The gamma distribution, as specified in our hierarchical model, is frequently used in models that include time delays in the production of mature functional proteins [Calderazzo *et al*., 2019; Krzyzanski, 2019; Tokuda *et al*., 2019; Korsbo and Jönsson, 2020]. To show that this assumption does not strongly bias the estimates, we used our model to infer delay parameters when the delay distribution was misspecified: With trajectories generated using beta and inverse-gamma delays our algorithm produced both population and individual estimates which were overall accurate, but still tended to slightly overestimate individual delay variances (see Fig. S8), as noted previously (Fig. 3b).

### 3.4 Estimation of time delay in transcriptional and translational regulation

We next used the hierarchical model with distributed delays to characterize the variability of delays and production rates in an experimentally observed clonal population. A birth-death process with distributed birth delays can be used to approximate the fluorescent protein (FP) production within individual cells: As with other proteins, the production of mature FP is not instantaneous. Cell growth does not strongly affect the production of FP [Austin *et al*., 2006; Megerle *et al*., 2008; Fritz *et al*., 2014], but is the main contributor to FP dilution, when FP is not enzymatically degraded [Andersen *et al*., 1998]. We assumed that protein expression is delayed but Poissonian. Transcriptional bursting, which can shape protein expression noise [Shahrezaei and Swain, 2008], can be modeled within our framework by including an additional transition processes which determines when transcription is “ON”.

We tested the performance of our hierarchical inference approach with data obtained using time– lapse fluorescent microscopy of a population of *E. coli* engineered to express a FP upon induction [Cheng *et al*. (2017)]. The population was observed in a microfluidic device, allowing for the recording of the flourescence signal at 1 min intervals from 39 cells and 27 cells in two independent experiments (Fig. **S4**a). The addition of Arabinose to the media at time *t* = 0 induced the transcription of yellow fluorescent protein (YFP) within all cells. As described previously, we assumed that the recorded fluorescent signal was proportional to the number of mature YFP molecules, thus allowing us to estimate the delay in the formation of the mature, fluorescing proteins after induction (For details see [Cheng *et al*., 2017; Choi *et al*., 2020)]. Previously, we performed a similar analysis assuming that cells in the population are identical. As noted previously, this assumption allowed us to increase power by pooling data across cells, but could lead to biases in population estimates, and did not allow us to estimate the variability in reaction rates and delays across the population.

**Fig. S4:**
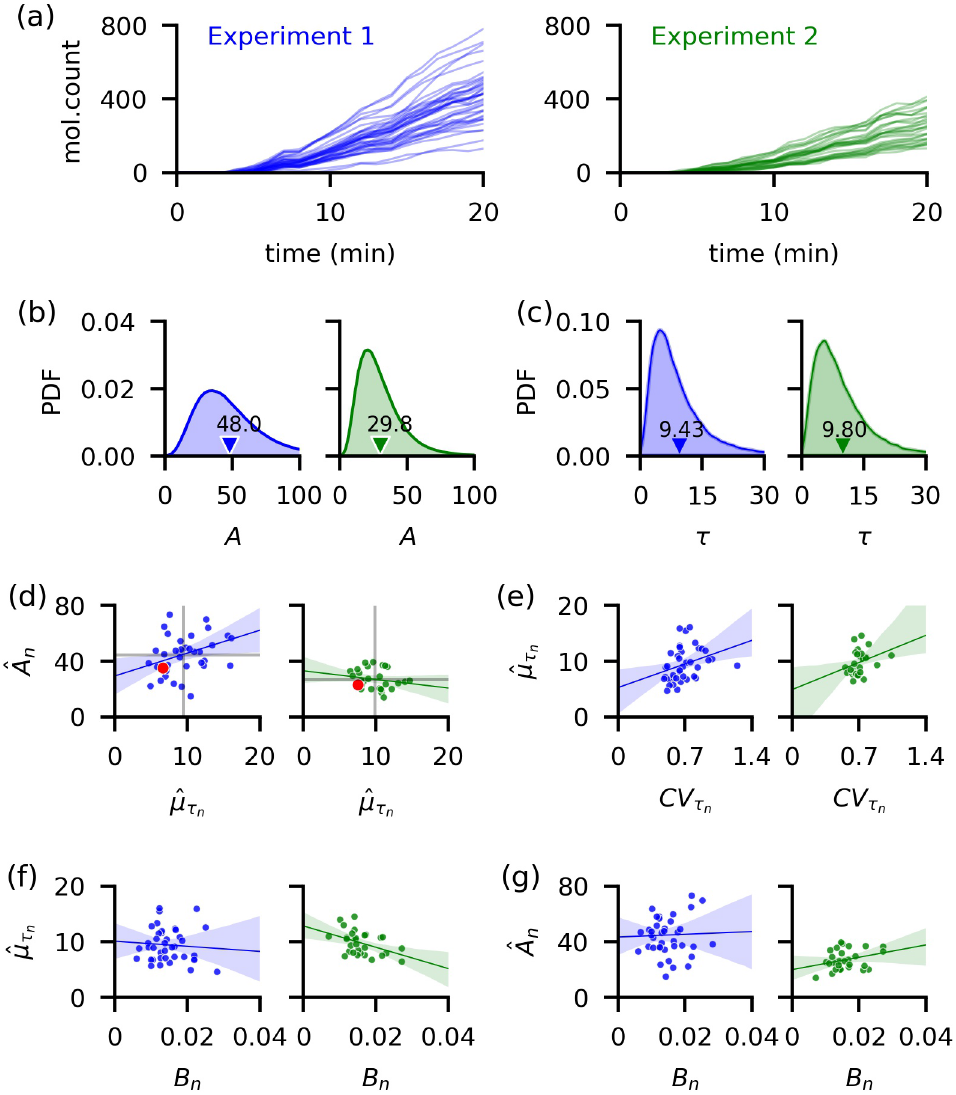
Consistent estimation of time delay distribution of YFP synthesis after induction. We use data from time-lapse images of YFP expression from two independent experiments performed previously by Cheng et al. [2017]. (a) Trajectory of estimated YFP molecule number obtained by dividing the total fluorescence level of each cell by a conversion constant. (b-c) We estimated the production rates, *A*_*n*_, and mean delay times, 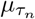, for each cell as the mean of the individual posterior distributions, obtained by fixing the dilution rate *B*_*n*_ estimated previously [Choi *et al*., 2020]. Because the molecular counts in the first were higher than in the second experiment, the population posterior mean for *A* was higher for the first. The population mean of the delay distributions are similar in the two experiments (9.43 and 9.80 min, respectively). (d) Both the average of the production rates and mean delay times (gray lines) are higher than previously reported by Choi et al. (red dots). We found no consistent correlation between *Â*_*n*_ and 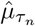 (*ρ* = 0.33 and *ρ* = −0.17) in the two experiments. (e) Individual *CV* s and 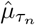 are moderately positively correlated (*ρ* = 0.31 and *ρ* = 0.30 in the first and second experiment, respectively). (f) The dilution rate, *B*_*n*_, and 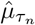 have *ρ* equal to −0.08 and −0.43 in the two experiments, respectively. (g) The reaction rates *B*_*n*_ and *A*_*n*_, show no clear evidence of correlation with *ρ* = −0.03 and *ρ* = 0.30 in the first and second experiment, respectively. Shaded regions in panels d-g show the 95% confidence interval for the regression estimate.

Since YFP did not saturate over the course of either experiment (Fig. **S4**a), we estimated the individual dilution rates *B*_*n*_, separately, and used the hierarchical model to estimate the individual production rates, *A*_*n*_, and delays, *τ*_*n*_. We used the dilution rate estimates by Choi et al. [2020], measured by tracking the rate of cell growth and division.

Because of the difference in the measured fluorescence levels in the two experiments (Fig. **S4**a), the estimated production rates across the population were higher in the first experiment (Fig. **S4**b). Despite this difference in the estimated production rates in the two experiments, the estimated mean delay times averaged across the population were close: 9.43 and 9.80 min (Fig. **S4**c). These delay estimates were about 3 min longer (Fig. **S4**d) than reported previously [Choi *et al*., 2020]. The estimated mean production rates were also somewhat higher than previously reported [Choi *et al*., 2020]. To check whether this discrepancy is due to a difference in the inference methods, we applied the two approaches to simulated trajectories obtained from populations of varying heterogeneity. While the hierarchical model produced good parameter estimates of individual cell parameters, a model that assumed that cells are identical lead to an underestimate of both the population birth rate and average delay that is similar to the discrepancy we observed here with experimental data, when cell-to-cell variability was high (See Fig. S6). Generating an ensemble of trajectories using the delay Gillespie algorithm and parameters sampled from the posterior distributions over the parameters for each individual cell produced good matches with experimental data (See Fig. S9).

We measured the distance between the population posterior densities of both the production rate, *A*, and delay time, *τ*, from the two experiments, by estimating the Kullback-Liebler (KL) divergence from the posterior samples [Wang *et al*., 2009]. Because of the considerable difference in YFP levels (Fig. **S4**a) in the two experiments, the KL divergence between the production rate posterior distributions (from the first experiment to the second) is large at 0.43. The posterior delay times (Fig. **S4**c), on the other hand, are almost identical with a KL divergence of 0.005. We also found that at the population level, the birth rate, *A*, has coefficients of variation (*CV* s) of 0.52 and 0.55 in the first and second experiment, respectively, while the collection of estimated mean delays, 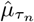 (Fig. **S4**d), has *CV* s of 0.31 and 0.21, respectively. This is similar to the *CV* of 0.20 reported for mean maturation times of YFP measured directly using fluorescent microscopy [Cheng *et al*., 2017].

To test for dependence between different parameter pairings we computed the Pearson correlation coefficients (Fig. **S4**d-g). We did not find any consistent relationships between the parameters, except for a moderate positive correlation between *CV* s and mean delay times, 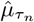, of the individual delay distribution (Fig. **S4**e).

The models we considered provide a minimal description of the underlying biological processes with delays. We therefore do not expect these models to overfit the data, and our tests indicate this is true. Cross validation and overfitting detection (see Fig. S10, S11, S12, S13) show that our hierarchical model with distributed delay extrapolates well to unobserved data, and is largely insensitive to small changes in input data, while the fixed delay model shows high bias. Using the distributed delay model to infer the full parameter set (i.e. including *B*_*n*_) from the initial portion of the data, produced results that were sensitive to input data resulting in dramatically different parameter estimates for different data sets, as well as unrealistically large parameter estimates that still produced good fits to the data. This indicated that parameters may not be recovered, and that the model can overfit the data when all parameters need to be inferred [Lever *et al*., 2016].

## 4 Conclusion

We have developed a hierarchical Bayesian framework for the inference of individual parameters of delayed biochemical reactions, as well as the distributions of these parameters across a population of observed reactions. We have shown that this method provides robust and accurate estimates of reaction rates, delay parameters, and their population distributions using both synthetic data and experimental measurements of gene regulatory networks obtained via fluorescent microscopy.

We demonstrated the performance and limitations of our method using realizations of a simple delayed birth-death reaction network. Although this is a simple setting, it shows that our algorithm can be used to simultaneously infer the dozens of parameters that characterize all the individuals in a population. Inferred reaction rate constants and delay distribution parameters, especially from experimental data, however, should be interpreted with care since our model does not take into account cell cycle descriptions in the dilution process [Beentjes *et al*., 2020]. Regulated birth and death reactions form the basis for more complex biochemical reaction networks. Thus, our method of deriving the likelihoods and approximate posteriors over the parameters, and the corresponding sampling algorithm are readily extendable to more complex systems with more species and reactions. However, the increased computational cost of evaluating more complex likelihoods, and sampling in high dimension will require alternate implementations of the sampling algorithm such as Hamiltonian Monte Carlo [McKay, 2003; Neal, 2011], variational approaches [Jordan *et al*., 1999; Wainwright *et al*., 2008], or machine learning methods [Cranmer *et al*., 2020].

We evaluated the performance of our algorithm assuming both fixed and distributed birth delays. The fixed delay model has fewer parameters, is easier to implement, faster to run, and performs better when measurements are generated using fixed delay processes. However, using the fixed delay model for inference leads to underestimates of birth reaction rates and delay times at the individual and population levels when reaction delays fluctuate. The hierarchical distributed delay model, on the other hand, is applicable more widely. It works well with high-resolution data, but at the expense of high computational cost. We also showed when data is sparsely sampled, the ensemble estimation allows our hierarchical model to outperform its non-hierarchical counterpart.

Our inference model is applicable to experimental data that can be obtained using fluorescent microscopy. In such experiments, cells can vary in growth rates, size, plasmid copy number, and other factors [Altschuler *et al*., 2010; Rinott *et al*., 2011; Snijder *et al*., 2011; Sherman *et al*., 2016; Mitchell *et al*., 2018]. Hierarchical models explicitly account for such heterogeneity and thus provide a suitable framework for inferring the variability and covariability of biochemical rates and delays across the population [Heydari *et al*., 2016; Tonner *et al*., 2020; Zechner *et al*., 2014]. Our estimates of the reaction rates and delays were higher than previous estimates [Choi *et al*., 2020]. This discrepancy may be due to biases introduced by data pooling without considering cell heterogeneity, as in the case of model implementation with synthetic data. Despite this difference in the parameter estimates, we were able to closely recreate the observed YFP molecular counts for all the cells using the resulting values from our hierarchical approach. Thus, we expect that our robust hierarchical approach can be extended to quantify the variabilty and covariabilty of complex gene network dynamics from experimental data.

