## Supplementary for "Hierarchical Bayesian models of transcriptional and translational regulation processes with delays"

### Supplementary Methods

Consider $N$ distinct observations of the same process $\mathbf{y}$. For every individual $n$, $n = 1, 2, \ldots, N$, we observe the evolution of $u$ species through $v$ reactions. Every reaction $R_{nk}$, $k = 1, 2, \ldots, v$, is endowed with a rate constant $\theta_{nk}$, so that reactions corresponding to individual $n$ are described by the rate vector $\theta_n = (\theta_{n1}, \theta_{n2}, \ldots, \theta_{nk}, \ldots, \theta_{nv})$. For a reaction $R_{nk}$ with delayed completion, we associate the delay parameters $\Delta_{nk} = (\Delta_{nk1}, \Delta_{nk2}, \ldots, \Delta_{nkl_k})$. The set of all delay parameters for an individual $n$ is written as $\Delta_n = \{\Delta_{nk}\}$. We denote by $\boldsymbol{\theta}$ the collection $\{\theta_n\}$ of all rate constants, and by $\boldsymbol{\Delta}$ the collection $\{\Delta_n\}$ of parameters that define all delay measures. For every individual observation, $\mathbf{y_n}$, we denote its subset of discrete-time observations as $\mathbf{y_{d,n}} = (y_n(0), y_n(1), \ldots, y_n(T-1), y_n(T))$.

Individual-level parameters $\theta_{nk}$ and $\Delta_{nkl}$ follow underlying distributions which are themselves characterized by hyperparameters, $\omega_{\theta_k}$ and $\omega_{\Delta_{kl}}$, respectively. As we specified gamma priors for each $\theta_{nk}$, the corresponding hyperparameter set for reaction $k$ is given by $\omega_{\theta_k} = \{a_{\theta_k}, b_{\theta_k}\}$. We denote the collection $\{\omega_{\theta_k}\}$ and $\{\omega_{\Delta_{kl}}\}$ of rate and delay hyperparameters respectively as $\omega_\theta$ and $\omega_\Delta$.

The MCMC algorithm to produce samples from the approximate posterior distribution obtained using the hierarchical model given by Eq. (6) can thus be described by the following steps.

1. For each $n = 1, 2, \ldots, N$, $k = 1, 2, \ldots, v$, and $i = 0, 1, \ldots, T-1$, initialize the number of reactions $r_{nki}$. Initialize the parameters $\boldsymbol{\theta}$ and $\boldsymbol{\Delta}$, and hyperparameters $\omega_\theta$ and $\omega_\Delta$.

2. For each $n$,

   (a) Sample, in order, $\theta_{nk}$, $k = 1, 2, \ldots, v$, given all rate hyperparameters $\omega_\theta$, other rate constants $\theta_{nm}$, $m \neq k$, delay parameters $\Delta_n$, and reaction numbers. If $y_n(t)$ and $\theta_{nk}$ are separable in $h_k(y_n(t), \theta_{nk})$, then sample $\theta_{nk}$ from the conjugate gamma distribution. Otherwise, use the Metropolis-Hastings algorithm.

   (b) Sample, in order, $\Delta_{nkl}$, $k = 1, 2, \ldots, v$ and $l = 1, 2, \ldots, l_k$, given all delay hyperparameters $\omega_\Delta$, other delay constants $\Delta_{nk'l'}$, $(k', l') \neq (k, l)$, rate parameters $\theta_n$, and reaction numbers, using the Metropolis-Hastings algorithm.

   (c) Update $r_{nki}$ for $k = 1, 2, ...v$ and $i = 0, 1, ..., T-1$, given $\theta_n$, $\Delta_n$, and the observed trajectory $y_n$ using the simplified block-updating method.

3. For every reaction $k$,

   (a) Sample $a_{\theta_k}$, given the rate constants $\{\theta_{nk}\}_n$ from the entire population and the other rate hyperparameter $b_{\theta_k}$.

   (b) Sample $b_{\theta_k}$, given the rate constants $\{\theta_{nk}\}_n$ from the entire population and other rate hyperparameter $a_{\theta_k}$.

   (c) Sample, in order, $\omega_{\Delta_{kls}}$, $l = 1, 2, \ldots, l_k$, $s = 1, 2, \ldots, |\omega_{\Delta_{kl}}|$ given the delay parameters $\{\Delta_{nkl}\}_n$ from the entire population and other delay hyperparameters $\omega_{\Delta_{kls'}}$, $s' \neq s$.

4. Repeat steps 2-3 until convergence.

Depending on the choice of hyperpriors, step 3 of this algorithm is either carried out by sampling from conjugate conditional distributions or using the Metropolis-Hastings algorithm. We provide below all likelihoods and resulting posterior distributions given specific hyperprior distributions and delay measures for a stochastic birth-death process with birth delays, dealt with in this work.

### Description of Bayesian inference of hierarchical model for a stochastic birth-death process with distributed birth delays

For the birth-death process, each individual $n$ has birth ($k = 1$) parameter $A_n$ and death ($k = 2$) parameter $B_n$, so that $\boldsymbol{\theta} = \{A_n, B_n\}_{n=1}^{N}$. We assumed that the completion of a birth reaction is delayed by a time $\tau_n$ following a gamma distribution $\Gamma(\alpha_n, \beta_n)$, so that $\boldsymbol{\Delta} = \{\alpha_n, \beta_n\}_{n=1}^{N}$. Since we only consider delays in the birth reaction, henceforth we write $\eta_n$ for the delay distribution $\eta_{n,1} = \Gamma(\tau_n; \alpha_n, \beta_n)$, and we write $\tau_n$ for $\tau_{n,1}$. With mass-action kinetics, the reaction hazards are given by

$$h_1(y_n(t), A_n) = A_n,$$
$$h_2(y_n(t), B_n) = B_n y_n(t).$$

In our setup where only discrete-time observations are available, only the birth reaction is delayed so that the corresponding average completion propensity for a birth reaction on the interval $(i, i+1]$ is

$$\hat{f}_1(i, \mathbf{y_{d,n}}, A_n, \Delta_n) = A_n \int_i^{i+1} \int_0^t d\eta(s) dt = A_n \int_i^{i+1} \frac{\gamma\left(\alpha_n, \beta_n \hat{t}\right)}{\Gamma\left(\alpha_n\right)} d\hat{t}, \tag{S1}$$

where $\Delta_n = \{\alpha_n, \beta_n\}$ and $\gamma\left(\alpha_n, \beta_n \hat{t}\right)$ is the lower Gamma incomplete function [Abramowitz and Stegun, 1965]. On the other hand, the death reaction propensity is given by

$$\hat{f}_2(i, \mathbf{y_{d,n}}, B_n) = \frac{h_2(y_n(i), B_n) + h_2(y_n(i+1), B_n)}{2} = \frac{B_n\left(y_n(i+1) + y_n(i)\right)}{2}, \tag{S2}$$

which is the average of the delay-free death reaction hazard between the times $i$ and $i+1$.

We use the approximate propensities Eq. (S1) and (S2) to define the total likelihood which accounts for $N$ individual trajectories, $\mathbf{y_d} = \{\mathbf{y_{d,n}}\}_n$, given by

$$\hat{\mathcal{L}}\left(\mathbf{y_d} \,|\boldsymbol{\theta}\,, \boldsymbol{\Delta}\right) = \prod_{n=1}^{N} \hat{L}\left(\mathbf{y_{d,n}} \,|\theta_n\,, \Delta_n\right), \tag{S3}$$

where

$$\hat{L}\left(\mathbf{y_{d,n}} \,|\theta_n\,, \Delta_n\right) = \prod_{i=0}^{T-1} \frac{\hat{f}_1(i, \mathbf{y_{d,n}}, A_n, \Delta_n)^{r_{n1i}}}{r_{n1i}!} \exp\left(-\hat{f}_1\left(i, \mathbf{y_{d,n}}, A_n, \Delta_n\right)\right)$$
$$\times \prod_{i=0}^{T-1} \frac{\hat{f}_2(i, \mathbf{y_{d,n}}, B_n)^{r_{n2i}}}{r_{n2i}!} \exp\left(-\hat{f}_2\left(i, \mathbf{y_{d,n}}, B_n\right)\right)$$

and $r_{nki}$, for $k = 1, 2$, is the number of reactions which completed in the time interval $(i, i+1]$.

Following the generative model shown in Fig. 2b, we specify gamma priors $\Gamma(A_n|a_A, b_A)$, $\Gamma(B_n|a_B, b_B)$, $\Gamma(\alpha_n|a_\alpha, b_\alpha)$, and $\Gamma(\beta_n|a_\beta, b_\beta)$ for $n = 1, ..., N$. For the reaction rate hyperparameters, we specified the improper joint hyperpriors $\pi(a_A, b_A) \propto \frac{1}{b_A}$ and $\pi(a_B, b_B) \propto \frac{1}{b_B}$. Denote the arbitrary hyperpriors $\pi(a_\alpha, b_\alpha)$ and $\pi(a_\beta, b_\beta)$ of $\alpha$ and $\beta$ respectively. We denote the collection, $\{a_A, a_B, b_A, b_B\}$, of reaction rate hyperparameters as $\omega_\theta$, and the collection of delay hyperparameters, $\{a_\alpha, a_\beta, b_\alpha, b_\beta\}$, as $\omega_\Delta$. Accounting for Eq.

(S1), (S2), and (S3), the joint posterior distribution over the parameters and hyperparameters is given by

$$\begin{aligned}
\pi\left(\boldsymbol{\theta}, \boldsymbol{\Delta}, \omega_\theta, \omega_\Delta \left| \mathbf{y_d} \right.\right) &\propto \pi\left(a_A, b_A\right)\pi\left(a_B, b_B\right)\pi\left(a_\alpha, b_\alpha\right)\pi\left(a_\beta, b_\beta\right)\hat{\mathcal{L}}\left(\mathbf{y_d}\left|\boldsymbol{\theta}, \boldsymbol{\Delta}\right.\right) \\
&\quad \times \prod_{n=1}^{N} \pi\left(A_n \left| a_A\right., b_A\right)\pi\left(B_n \left| a_B\right., b_B\right)\pi\left(\alpha_n \left| a_\alpha\right., b_\alpha\right)\pi\left(\beta_n \left| a_\beta\right., b_\beta\right) \\
&= \frac{1}{b_A}\frac{1}{b_B}\pi\left(a_\alpha, b_\alpha\right)\pi\left(a_\beta, b_\beta\right) \\
&\quad \times \prod_{n=1}^{N}\prod_{i=0}^{T-1} \frac{\left(A_n \int_i^{i+1} \frac{\gamma\left(\alpha_n, \beta_n \hat{t}\right)}{\Gamma(\alpha_n)} d\hat{t}\right)^{r_{n1i}}}{r_{n1i}!} \exp\left(-A_n \int_i^{i+1} \frac{\gamma\left(\alpha_n, \beta_n \hat{t}\right)}{\Gamma\left(\alpha_n\right)} d\hat{t}\right) \\
&\quad \times \prod_{n=1}^{N}\prod_{i=0}^{T-1} \frac{\left[(1/2)B_n\left(y_n\left(i+1\right)+y_n\left(i\right)\right)\right]^{r_{n2i}}}{r_{n2i}!} \exp\left(-(1/2)B_n\left(y_n\left(i+1\right)+y_n\left(i\right)\right)\right) \\
&\quad \times \prod_{n=1}^{N} \frac{{b_A}^{a_A}}{\Gamma\left(a_A\right)} {A_n}^{a_A - 1}\exp\left(-b_A A_n\right) \frac{{b_B}^{a_B}}{\Gamma\left(a_B\right)} {B_n}^{a_B - 1}\exp\left(-b_B B_n\right) \\
&\quad \times \prod_{n=1}^{N} \frac{{b_\alpha}^{a_\alpha}}{\Gamma\left(a_\alpha\right)} {\alpha_n}^{a_\alpha - 1}\exp\left(-b_\alpha \alpha_n\right) \frac{{b_\beta}^{a_\beta}}{\Gamma\left(a_\beta\right)} {\beta_n}^{a_\beta - 1}\exp\left(-b_\beta \beta_n\right).
\end{aligned} \tag{S4}$$

Without specifying hyperpriors for the delay parameters $\boldsymbol{\Delta}$, using Eq. (S4), we can derive the conditional posterior of $A_n$ and $B_n$, which belong to the Gamma family:

$$\begin{aligned}
A_n \left| y, a_A, b_A, \Delta_n \right. &\sim \Gamma\left(\sum_{i=0}^{T-1} r_{n1i} + a_A, \sum_{i=0}^{T-1} \int_i^{i+1} \frac{\gamma\left(\alpha_n, \beta_n \hat{t}\right)}{\Gamma\left(\alpha_n\right)} d\hat{t} + b_A\right), \\
B_n | y, a_B, b_B &\sim \Gamma\left(\sum_{i=0}^{T-1} r_{n2i} + a_B\,, \sum_{i=0}^{T-1} \frac{y_n(i+1)+y_n(i)}{2} + b_B\right).
\end{aligned} \tag{S5}$$

The delay parameters $\alpha_n$ and $\beta_n$ do not have standard distributions as conditional posteriors which are proportional to

$$\begin{aligned}
\alpha_n \left| \mathbf{y_{d,n}}, A_n, \beta_n \right. &\propto \prod_{i=0}^{T-1} \left[\int_i^{i+1} \frac{\gamma\left(\alpha_n, \beta_n \hat{t}\right)}{\Gamma\left(\alpha_n\right)} d\hat{t}\right]^{r_{n1i}} \exp\left(-A_n \sum_{i=0}^{T-1} \int_i^{i+1} \frac{\gamma\left(\alpha_n, \beta_n \hat{t}\right)}{\Gamma\left(\alpha_n\right)} d\hat{t}\right) {\alpha_n}^{a_\alpha - 1}\exp\left(-\alpha_n b_\alpha\right), \\
\beta_n \left| \mathbf{y_{d,n}}, A_n, \alpha_n \right. &\propto \prod_{i=0}^{T-1} \left[\int_i^{i+1} \frac{\gamma\left(\alpha_n, \beta_n \hat{t}\right)}{\Gamma\left(\alpha_n\right)} d\hat{t}\right]^{r_{n1i}} \exp\left(-A_n \sum_{i=0}^{T-1} \int_i^{i+1} \frac{\gamma\left(\alpha_n, \beta_n \hat{t}\right)}{\Gamma\left(\alpha_n\right)} d\hat{t}\right) {\beta_n}^{a_\beta - 1}\exp\left(-\beta_n b_\beta\right).
\end{aligned} \tag{S6}$$

The shape parameters of the hyperpriors for the reaction rate constants $A$ and $B$ do not have conditional posteriors which are known distributions but are proportional to:

$$\begin{aligned}
\pi\left(a_A \left| A, b_A \right.\right) &\propto \frac{b_A^{N a_A}}{\Gamma(a_A)^N} \prod_{n=1}^{N} A_n^{a_A - 1}, \\
\pi\left(a_B \left| B, b_B \right.\right) &\propto \frac{b_B^{N a_B}}{\Gamma(a_B)^N} \prod_{n=1}^{N} B_n^{a_B - 1},
\end{aligned} \tag{S7}$$

while the rate parameters of the hyperpriors for $A$ and $B$ belong to the gamma family:

$$\begin{aligned} b_A \,|A, a_A &\sim \Gamma\left(Na_A, \sum_{n=1}^{N} A_n\right), \\ b_B \,|B, a_B &\sim \Gamma\left(Na_B, \sum_{n=1}^{N} B_n\right). \end{aligned} \tag{S8}$$

The choice of hyperpriors for the delay hyperparameters dictates what the conditional posterior distributions of $a_\alpha$, $a_\beta$, $b_\alpha$, and $b_\beta$ will be. We present derivations using three different choices of delay hyperprior distributions. We first show the cases of the non-informative rational hyperprior and maximal data information prior (MDIP), and afterwards the informative folded normal distribution.

A typical non-informative joint hyperprior is the rational prior, which for the pair $(a, b)$ takes the form $\pi(a, b) = \frac{1}{b}$. Setting such hyperpriors for the hyperparameters corresponding to both $\alpha$ and $\beta$ yields conjugate conditional posteriors for $b_\alpha$ and $b_\beta$ that belong to the Gamma family. This choice of hyperprior, however, is not conjugate for both $a_\alpha$ and $a_\beta$. The conditional posteriors for the hyperparameters are given by

$$\begin{aligned} \pi\left(a_\alpha \,|\alpha, b_\alpha\right) &\propto \frac{b_\alpha^{Na_\alpha}}{\Gamma(a_\alpha)^N} \prod_{n=1}^{N} \alpha_n^{a_\alpha - 1}, \\ \pi\left(a_\beta \,|\beta, b_\beta\right) &\propto \frac{b_\beta^{Na_\beta}}{\Gamma(a_\beta)^N} \prod_{n=1}^{N} \beta_n^{a_\beta - 1}, \\ b_\alpha \,|\alpha, a_\alpha &\sim \Gamma\left(Na_\alpha, \sum_{n=1}^{N} \alpha_n\right), \\ b_\beta \,|\beta, a_\beta &\sim \Gamma\left(Na_\beta, \sum_{n=1}^{N} \beta_n\right). \end{aligned} \tag{S9}$$

The maximal data information prior (MDIP) [Pradhan *et al.*, 2011; Zellner, 1991] is derived by maximizing the Kullback-Leibler divergence between the data density and the prior distribution. In our generative model, each of the parameters of an individual delay distribution is sampled from a gamma distribution $\Gamma(x; a, b)$ thereby serving as prior distribution in the hierarchical inference. In this case the MDIP for the hyperparameters $(a, b)$ becomes

$$\pi\left(a, b\right) = \frac{b}{\Gamma\left(a\right)} \exp\left\{\left(a - 1\right) \frac{\psi\left(a\right)}{\Gamma(a)} - a\right\}, \tag{S10}$$

where $\psi(a) = \frac{\Gamma'(a)}{\Gamma(a)}$ is the digamma function [Moala *et al.*, 2013].

The resulting conditional posterior for $a_\alpha$ and $a_\beta$ do not follow known distributions, however the MDIP is a conjugate prior for both $b_\alpha$ and $b_\beta$ whose conditional posteriors belong to the gamma family. The conditional posteriors for the hyperparameters are given by

$$\begin{aligned}
\pi\left(a_\alpha \,|\alpha, b_\alpha\right) &\propto \frac{b_\alpha^{Na_\alpha}}{\Gamma(a_\alpha)^{N+1}} \prod_{n=1}^{N} \alpha_n^{a_\alpha-1} \exp\left\{(a_\alpha-1)\,\frac{\psi\left(a_\alpha\right)}{\Gamma(a_\alpha)} - a_\alpha\right\}, \\
\pi\left(a_\beta \,|\beta, b_\beta\right) &\propto \frac{b_\beta^{Na_\beta}}{\Gamma(a_\beta)^{N+1}} \prod_{n=1}^{N} \beta_n^{a_\beta-1} \exp\left\{(a_\beta-1)\,\frac{\psi\left(a_\beta\right)}{\Gamma(a_\beta)} - a_\beta\right\}, \\
b_\alpha \,|\alpha, a_\alpha &\sim \Gamma\left(Na_\alpha + 2, \sum_{n=1}^{N} \alpha_n\right), \\
b_\beta \,|\beta, a_\beta &\sim \Gamma\left(Na_\beta + 2, \sum_{n=1}^{N} \beta_n\right).
\end{aligned} \tag{S11}$$

Since the delay hyperparameters are positive, being parameters of a Gamma distribution, the joint folded normal distribution [Leone *et al.*, 1961; Psarakis *et al.*, 2001] is a candidate hyperprior distribution that can effectively define an arbitrarily strong joint hyperprior for these hyperparameters. The bivariate version follows naturally from the bivariate Gaussian distribution which describes two non-negative real-valued random variables $X$ and $Y$ with probability density function given by

$$\begin{aligned}
f\left(x, y\right) =& \frac{1}{2\pi\sigma_1\sigma_2\sqrt{1-\rho^2}} \\
&\times \left\{ \exp\left(-\frac{1}{2\left(1-\rho^2\right)}\left(\frac{\left(x-\mu_1\right)^2}{{\sigma_1}^2} - 2\rho\frac{\left(y-\mu_1\right)\left(x-\mu_2\right)}{\sigma_1\sigma_2} + \frac{\left(y-\mu_2\right)^2}{{\sigma_2}^2}\right)\right)\right. \\
&+ \exp\left(-\frac{1}{2\left(1-\rho^2\right)}\left(\frac{\left(x+\mu_1\right)^2}{{\sigma_1}^2} - 2\rho\frac{\left(x+\mu_1\right)\left(x+\mu_2\right)}{\sigma_1\sigma_2} + \frac{\left(y+\mu_2\right)^2}{{\sigma_2}^2}\right)\right) \\
&+ \exp\left(-\frac{1}{2\left(1-\rho^2\right)}\left(\frac{\left(x+\mu_1\right)^2}{{\sigma_1}^2} + 2\rho\frac{\left(x+\mu_1\right)\left(y-\mu_2\right)}{\sigma_1\sigma_2} + \frac{\left(y-\mu_2\right)^2}{{\sigma_2}^2}\right)\right) \\
&\left.+ \exp\left(-\frac{1}{2\left(1-\rho^2\right)}\left(\frac{\left(x-\mu_1\right)^2}{{\sigma_1}^2} + 2\rho\frac{\left(x-\mu_1\right)\left(y+\mu_2\right)}{\sigma_1\sigma_2} + \frac{\left(y+\mu_2\right)^2}{{\sigma_2}^2}\right)\right)\right\},
\end{aligned}$$

where $x > 0$, $y > 0$, $\sigma_i > 0$, $\mu_i \in \mathbb{R}$, $i = 1, 2$, and $|\rho| \leq 1$.

This distribution is not conjugate for any of the delay hyperparameters and the conditional posterior distributions resulting from this choice are given by

$$\begin{aligned}
\pi\left(a_\alpha \,|\{\alpha_n\}_n, b_\alpha\right) &\propto \frac{b_\alpha^{Na_\alpha}}{\Gamma(a_\alpha)^{N}} \prod_{n=1}^{N} \alpha_n^{a_\alpha-1} f(a_\alpha, b_\alpha; \mu_{a_\alpha}, \sigma_{a_\alpha}, \mu_{b_\alpha}, \sigma_{b_\alpha}, \rho_\alpha), \\
\pi\left(a_\beta \,|\{\beta_n\}_n, b_\beta\right) &\propto \frac{b_\beta^{Na_\beta}}{\Gamma(a_\beta)^{N}} \prod_{n=1}^{N} \beta_n^{a_\beta-1} f(a_\beta, b_\beta; \mu_{a_\beta}, \sigma_{a_\beta}, \mu_{b_\beta}, \sigma_{b_\beta}, \rho_\beta), \\
\pi\left(b_\alpha \,|\{\alpha_n\}_n, a_\alpha\right) &\propto b_\alpha^{Na_\alpha} \exp\left(-b_\alpha \sum_{n=1}^{N} \alpha_n\right) f(a_\alpha, b_\alpha; \mu_{a_\alpha}, \sigma_{a_\alpha}, \mu_{b_\alpha}, \sigma_{b_\alpha}, \rho_\alpha), \\
\pi\left(b_\beta \,|\{\beta_n\}_n, a_\beta\right) &\propto b_\beta^{Na_\beta} \exp\left(-b_\beta \sum_{n=1}^{N} \beta_n\right) f(a_\beta, b_\beta; \mu_{a_\beta}, \sigma_{a_\beta}, \mu_{b_\beta}, \sigma_{b_\beta}, \rho_\beta),
\end{aligned} \tag{S12}$$

where the corresponding folded normal hyperprior $f(a_Z, b_Z)$ is parametrized by $\mu_{a_Z}, \sigma_{a_Z}$, $\mu_{b_Z}, \sigma_{b_Z}, \rho_Z$ for $Z \in \{\alpha, \beta\}$. The amount of information about a delay parameter $Z$ is controlled by how close $\mu_{a_Z}$ and $\mu_{b_Z}$ are to the true values, and how small $\sigma_{a_Z}$ and $\sigma_{b_Z}$ are.

The MCMC algorithm for the stochastic birth-death process with distributed birth delays proceeds as follows.

1. For each $n$ and $i$, for $n = 1, 2, \ldots, N$ and $i = 0, 1, \ldots, T-1$, initialize the number of reactions by setting $r_{n1i} = y_n(i+1) - y(i)$ and $r_{n2i} = 0$ if $y_n(i+1) \geq y_n(i)$, otherwise $r_{n2i} = y_n(i) - y_n(i+1)$ and $r_{n1i} = 0$. Initialize $a_A, a_B, b_A, b_B$ using appropriate values. Initialize $A_n$ and $B_n$ by sampling from their conjugate gamma posterior distributions (Eq. (S5)), and set an appropriate values for $\alpha_n$ and $\beta_n$.

2. For each $n$,

   (a) Sample $A_n$ and $B_n$ from their conditional conjugate posterior distribution given by Eq. (S5).

   (b) Since the conditional posterior for $\alpha_n$ and $\beta_n$ do not follow a known distributions (Eq. (S6)), use the Metropolis-Hastings algorithm to draw samples, in order, from the conditional posterior $\alpha_n \left| \mathbf{y_{d,n}}, A_n, \beta_n \right.$ and $\beta_n \left| \mathbf{y_{d,n}}, A_n, \alpha_n \right.$. We used the truncated Gaussian distribution with positive support as proposal distribution for $\alpha_n$ and a gamma proposal for $\beta_n$ [Choi *et al.*, 2020].

   (c) Conditioned on $A_n$, $B_n$, $\alpha_n$, and $\beta_n$, for each time index $i$, update $r_{n1i}$ and $r_{n2i}$. As the number of reactions are not observed directly, we will sample over them by following a block-updating method [Boys *et al.*, 2008] which uses a random walk proposal on the number of birth reactions. We use the Metropolis-Hastings algorithm with a random walk chain to sample the number of completions of each type of reaction in a time interval $(i, i+1]$, $i = 0, 1, \ldots T-1$, given the observed system states $y_n(i)$ and $y_n(i+1)$. For the $i^{\text{th}}$ interval, the joint conditional posterior of $r_{n1i}$ and $r_{n2i}$ is given by

$$\pi\left(r_{n1i}, r_{n2i} \left| \mathbf{y_{d,n}}, A_n, B_n, \Delta_n \right.\right) \propto \frac{\left(A_n \int_i^{i+1} \frac{\gamma\left(a_n, \beta_n \hat{t}\right)}{\Gamma\left(\alpha_n\right)} d\hat{t}\right)^{r_{n1i}}}{r_{n1i}!} \frac{\left[B_n\left(y_n\left(i\right) + y_n\left(i+1\right)\right)/2\right]^{r_{n2i}}}{r_{n2i}!},$$

   which is proportional to the product of the density functions of Poisson random variables. Denote by $r_{n1i}^{(j-1)}$ the value of $r_{n1i}$ from the $(j-1)^{\text{th}}$ iteration. The proposal distribution can be chosen to be a discrete random walk in which the current value is augmented by $u$, that is the difference of two Poisson random variables whose means are both equal to some $\lambda$ which is usually a function of $r_{n1i}^{(j-1)}$. This value $u$ is then used to define the proposed value $r_{n1i}^{*} = r_{n1i}^{(j-1)} + u$. In particular, the distribution of the update value $u$ is a Skellam distribution [Boys *et al.*, 2008; Johnson *et al.*, 1969] given by

$$p\left(u \left| {r_{n1i}}^{(j-1)} \right.\right) = \exp\left(-2{r_{n1i}}^{(j-1)}\right) I_u\left(2{r_{n1i}}^{(j-1)}\right),$$

   where $I_u$ is a regular modified Bessel function of order $u$. Once $r_{n1i}^{*}$ is chosen, then $r_{n2i}^{*}$ can be uniquely determined using $y_n(i+1) - y_n(i) = r_{n1i}^{*} - r_{n2i}^{*}$.

3. Sample the hyperparameters which describe the distribution of $A_n$, $B_n$, $\alpha_n$, and $\beta_n$ across the population.

   (a) As the conditional posteriors (Eq. (S7)) of $a_A$ and $a_B$ are not known distributions, we draw samples using the Metropolis-Hastings algorithm. We specified as proposal distribution the truncated Gaussian distribution with positive support.

   (b) Sample $b_A$ and $b_B$ from their conditional conjugate posterior distributions given by Eq. (S8).

   (c) For rational priors, use Eq. (S9) to implement the Metropolis-Hasting algorithm with a positively-supported Gaussian proposal distribution to sample $a_\alpha$ and $a_\beta$ from their conditional posterior. Sample $b_\alpha$ and $b_\beta$ from their conjugate gamma conditional posteriors.
   In the case of the MDIP, use Eq. (S11) to implement the Metropolis-Hasting algorithm with a positively-supported Gaussian proposal distribution to sample $a_\alpha$ and $a_\beta$ from their conditional posterior. Sample $b_\alpha$ and $b_\beta$ from their conjugate gamma conditional posterior.
   If folded normal distributions are used as hyperprior for $\alpha_n$ and $\beta_n$, we use Eq. (S12), to implement the Metropolis-Hasting algorithm with a positively-supported Gaussian proposal distribution to sample $a_\alpha$, $a_\beta$, $b_\alpha$, and $b_\beta$ from their conditional posteriors.

4. Repeat steps 1-3 until convergence.

### Description of Bayesian inference of a hierarchical model for a stochastic birth-death process with fixed birth delays

Similar to the stochastic birth-death process with distributed delays, here each individual $n$ has birth ($k = 1$) parameter $A_n$ and death ($k = 2$) parameter $B_n$, so that $\boldsymbol{\theta} = \{A_n, B_n\}_{n=1}^{N}$. Since we are fixing the delays in each experiment, the delay measure $\eta_{n,k}$ is the Dirac point mass measure centered at the fixed delay $\tau_{n,k}$. Since we only consider delays in the birth reaction, henceforth we write $\eta_n$ for $\eta_{n,1}$ and we write $\tau_n$ for $\tau_{n,1}$. With mass-action kinetics, the reaction hazards are given by

$$h_1(y_n(t), A_n) = A_n,$$
$$h_2(y_n(t), B_n) = B_n y_n(t).$$

With only the discrete-time observations, the average completion propensity for a birth reaction on the interval $(i, i+1]$ is

$$\hat{f}_1(i, \mathbf{y_{d,n}}, A_n, \Delta_n) = A_n \int_i^{i+1} \int_0^t d\eta_n(s)dt = A_n \cdot p_{n,i}, \tag{S13}$$

where $\Delta_n = \{\tau_n\}$ and $p_{n,i} = \begin{cases} 0 & \text{if } i+1 \leqslant \tau_n \\ \min(1, i+1-\tau_n) & \text{otherwise} \end{cases}$. On the other hand, the average completion propensity for the death reaction is the same as Eq. (S2):

$$\hat{f}_2(i, \mathbf{y_{d,n}}, B_n) = \frac{h_2(y_n(i), B_n) + h_2(y_n(i+1), B_n)}{2} = \frac{B_n\left(y_n(i+1) + y_n(i)\right)}{2}.$$

Using Eq. (S13) and (S2), we obtain the total likelihood for $\mathbf{y_d} = \{\mathbf{y_{d,n}}\}_n$,

$$\hat{\mathcal{L}}\left(\mathbf{y_d} \,|\boldsymbol{\theta}\ , \boldsymbol{\Delta}\right) = \prod_{n=1}^{N} \hat{L}\left(\mathbf{y_{d,n}} \,|\theta_n\ , \Delta_n\right), \tag{S14}$$

where

$$\begin{aligned}\hat{L}\left(\mathbf{y_{d,n}} \,|\theta_n\ , \Delta_n\right) = &\prod_{i=0}^{T-1} \frac{\hat{f}_1(i, \mathbf{y_{d,n}}, A_n, \Delta_n)^{r_{n1i}}}{r_{n1i}!} \exp\left(-\hat{f}_1\left(i, \mathbf{y_{d,n}}, A_n, \Delta_n\right)\right) \\ &\times \prod_{i=0}^{T-1} \frac{\hat{f}_2(i, \mathbf{y_{d,n}}, B_n)^{r_{n2i}}}{r_{n2i}!} \exp\left(-\hat{f}_2\left(i, \mathbf{y_{d,n}}, B_n\right)\right)\end{aligned}$$

and $r_{nki}$, for $k = 1, 2$, is the number of reactions which completed in the time interval $(i, i+1]$.

Following the generative model (See Fig. 1a in main text), we use Gamma priors $\Gamma(A|a_A, b_A)$, $\Gamma(B|a_B, b_B)$, and $\Gamma(\tau|a_\tau, b_\tau)$ for $A_n$, $B_n$, and $\tau_n$ respectively for $n = 1, ..., N$. We also specified the improper joint hyperpriors $\pi(a_A, b_A) \propto \dfrac{1}{b_A}$, $\pi(a_B, b_B) \propto \dfrac{1}{b_B}$, and $\pi(a_\tau, b_\tau) \propto \dfrac{1}{b_\tau}$. We denote the collection, $\{a_A, a_B, b_A, b_B\}$, of reaction rate hyperparameters as $\omega_\theta$, and the collection of delay hyperparameters, $\{a_\tau, b_\tau\}$, as $\omega_\Delta$. Integrating all details from Eq. (S13), (S2), and (S14) we arrive at the total posterior distribution over the parameters and hyperparameters

$$\begin{aligned}
\pi\left(\boldsymbol{\theta},\boldsymbol{\Delta},\omega_\theta,\omega_\Delta\left|\mathbf{y_d}\right.\right) &\propto \pi\left(a_A,b_A\right)\pi\left(a_B,b_B\right)\pi\left(a_\tau,b_\tau\right)\hat{\mathcal{L}}\left(\mathbf{y_d}\left|\boldsymbol{\theta},\boldsymbol{\Delta}\right.\right)\\
&\quad\times\prod_{n=1}^{N}\pi\left(A_n\left|a_A\right.,b_A\right)\pi\left(B_n\left|a_B\right.,b_B\right)\pi\left(\tau_n\left|a_\tau\right.,b_\tau\right)\\
&=\frac{1}{b_A}\frac{1}{b_B}\frac{1}{b_\tau}\prod_{n=1}^{N}\prod_{i=0}^{T-1}\frac{\left(A_np_{n,i}\right)^{r_{n1i}}}{r_{n1i}!}\exp\left(-A_np_{n,i}\right)\\
&\quad\times\prod_{n=1}^{N}\prod_{i=0}^{T-1}\frac{\left[(1/2)B_n\left(y_n\left(i+1\right)+y_n\left(i\right)\right)\right]^{r_{n2i}}}{r_{n2i}!}\exp\left(-(1/2)B_n\left(y_n\left(i+1\right)+y_n\left(i\right)\right)\right)\\
&\quad\times\prod_{n=1}^{N}\frac{{b_A}^{a_A}}{\Gamma\left(a_A\right)}{A_n}^{a_A-1}\exp\left(-b_AA_n\right)\frac{{b_B}^{a_B}}{\Gamma\left(a_B\right)}{B_n}^{a_B-1}\exp\left(-b_BB_n\right)\frac{{b_\tau}^{a_\tau}}{\Gamma\left(a_\tau\right)}{\tau_n}^{a_\tau-1}\exp\left(-b_\tau\tau_n\right).
\end{aligned}\tag{S15}$$

Using Eq. (S15), we derive the conditional posterior distributions of the parameters and hyperparameters. For each $A_n$ and $B_n$, we obtain the conditional posteriors which belong to the gamma family:

$$\begin{aligned}
A_n\left|\mathbf{y_{d,n}}\right.,a_A,b_A,\tau_n &\sim \Gamma\left(\sum_{i=0}^{T-1}r_{n1i}+a_A,T-\tau_n+b_A\right),\\
B_n|\mathbf{y_{d,n}},a_B,b_B &\sim \Gamma\left(\sum_{i=0}^{T-1}r_{2ni}+a_B\ ,\ \sum_{i=0}^{T-1}\frac{y_n(i+1)+y_n(i)}{2}+b_B\right).
\end{aligned}\tag{S16}$$

The conditional posterior for a delay parameter, on the other hand, does not follow a known distribution and is proportional to:

$$\pi\left(\tau_n\left|\mathbf{y_{d,n}}\right.,a_\tau,b_\tau,A_n\right)\propto\left(\prod_{i=0}^{T-1}{p_{n,i}}^{r_{n1i}}\right)\exp\left(-A_n\left(T-\tau_n\right)\right){\tau_n}^{a_\tau-1}\exp\left(-b_\tau\tau_n\right).\tag{S17}$$

The shape parameters of hyperprior for the reaction constants $A$ and $B$ do not have known distribution as conditional posteriors and are proportional to:

$$\begin{aligned}
\pi(a_A|b_A,A) &\propto \frac{b_A^{Na_A}}{\Gamma(a_A)}\prod_{n=1}^{N}{A_n}^{a_A-1},\\
\pi(a_B|b_B,B) &\propto \frac{b_B^{Na_B}}{\Gamma(a_B)}\prod_{n=1}^{N}{B_n}^{a_B-1}.
\end{aligned}\tag{S18}$$

The rate parameters of hyperprior for the reaction constant $A$ and $B$ belong to the gamma family:

$$\begin{aligned}
b_A|a_A,A &\sim \Gamma(Na_A\ ,\ \sum_{n=1}^{N}A_n),\\
b_B|a_B,B &\sim \Gamma(Na_B\ ,\ \sum_{n=1}^{N}B_n).
\end{aligned}\tag{S19}$$

The conditional posterior for hyper parameters of delay follow the same form as the ones for the reaction rate constants.

$$\begin{aligned}
\pi(a_\tau|b_\tau,\tau) &\propto \frac{b_\tau^{Na_\tau}}{\Gamma(a_\tau)}\prod_{n=1}^{N}{\tau_n}^{a_\tau-1},\\
b_\tau|a_\tau,\tau &\sim \Gamma(Na_\tau\ ,\ \sum_{n=1}^{N}\tau_n).
\end{aligned}\tag{S20}$$

The MCMC algorithm for the hierarchical model of the stochastic birth-death process with constant birth delays proceeds as follows.

1. For each $n$ and $i$, for $n = 1, 2, \ldots, N$ and $i = 0, 1, \ldots, T-1$, initialize the number of reactions by setting $r_{n1i} = y_n(i+1) - y_n(i)$ and $r_{n2i} = 0$ if $y_n(i+1) \geq y_n(i)$, otherwise $r_{n2i} = y_n(i+1) - y(i)$ and $r_{n1i} = 0$. Initialize the hyperparameters $a_A, a_B, a_\tau, b_A, b_B, b_\tau$ using appropriate values, and initialize $A_n$ and $B_n$ by sampling from their conjugate gamma posterior distributions (Eq. (S16)). Set an appropriate value for $\tau_n$.

2. For each $n$,

   (a) Sample $A_n$ and $B_n$ from their conditional conjugate posterior distribution given by Eq. (S16).

   (b) Since the conditional posterior for $\tau_n$ does not follow a known distribution (Eq. (S17)), we used the Metropolis-Hastings algorithm to iteratively draw samples from the conditional posterior $\tau_n \,|\mathbf{y_{d,n}}, a_\tau, b_\tau, A_n$. We used the truncated Gaussian distribution with positive support as proposal distribution.

   (c) The update process for the number of completed reactions $r_{nki}$ is similar to the case of distributed birth delays, only that we will change the mean of the Poisson likelihood for the birth reaction to $\hat{f}_1(i, \mathbf{y_{d,n}}, A_n, \Delta_n) = A_n \cdot p_{n,i}$ where $\Delta_n = \{\tau_n\}$ and $p_{n,i} = \begin{cases} 0 & \text{if } i+1 \leqslant \tau_n \\ \min(1, i+1-\tau_n) & \text{otherwise} \end{cases}$. Hence for the interval $(i, i+1]$ interval, the joint conditional posterior of $r_{n1i}$ and $r_{n2i}$ is given by

$$\pi\left(r_{n1i}, r_{n2i} \,|\mathbf{y_{d,n}}, A_n, B_n, \Delta_n\right) \propto \frac{\left(A_n \cdot p_{n,i}\right)^{r_{n1i}}}{r_{n1i}!} \frac{\left[B_n\left(y_n(i) + y_n(i+1)\right)/2\right]^{r_{n2i}}}{r_{n2i}!}.$$

3. We now sample the hyperparameters which describe the distribution of $A_n$, $B_n$, and $\tau_n$ across the population.

   (a) As the conditional posteriors (Eq. (S18)) of $a_A$ and $a_B$ are not known distributions, we draw samples using the Metropolis-Hastings algorithm. We specified as proposal distribution the truncated Gaussian distribution with positive support.

   (b) Sample $b_A$ and $b_B$ from their conditional conjugate posterior distributions given by Eq. (S19).

   (c) With Eq. (S20), use the Metropolis-Hasting algorithm with a positively-supported Gaussian proposal to sample $a_\tau$ from its conditional posterior. Sample $b_\tau$ from its conjugate gamma conditional posterior.

4. Repeat steps 1-3 until convergence.

### Sampling from population distributions and individual delay distributions

Here we present the algorithm for generating the posterior population distributions and the individual delay distributions found throughout the main text. Population distributions are not directly sampled in the algorithm, and we instead sample from the posterior distribution of the hyperparameters. Here, we present the algorithm we used to sample from the population marginal posteriors of the parameters $A$, $\alpha$, $\beta$, and the delay time $\tau$. We also apply the same strategy to sample values from the individual delay distributions.

Through the MCMC algorthm we obtain the hyperparameter posterior distributions $\pi(a_Z)$ and $\pi(b_Z)$, for $Z \in \{A, B, \tau\}$ in the fixed delay, and $Z \in \{A, B, \alpha, \beta\}$ in the distributed delay case. We employed the algorithm below to sample from the respective population distributions.

1. Take $m$ samples $a_Z^s$ from $\pi(a_Z)$.

2. Take $m$ samples $b_Z^s$ from $\pi(b_Z)$.

3. For each pair $(a_Z^s, b_Z^s)$, take $p$ samples from $\Gamma(a_Z^s, b_Z^s)$.

4. Combine all the $mp$ samples taken from step 3. This pooled samples are realizations of the population distribution of the reaction rate or delay parameter $Z$.

In the distributed delay case, a similar algorithm was also applied to sample from the individual delay posterior distributions. For an individual $n$, the algorithm infers the posteriors $\pi(\alpha_n)$ and $\pi(\beta_n)$. We sample from $\pi(\tau_n)$ as follows.

1. Take $m$ samples $\alpha_n^s$ from $\pi(\alpha_n)$.
2. Take $m$ samples $\beta_n^s$ from $\pi(\beta_n)$.
3. For each pair $(\alpha_n^s, \beta_n^s)$, take $p$ samples from $\Gamma(\alpha_n^s, \beta_n^s)$.
4. Combine all the $mp$ samples taken from step 3. The pooled samples are realizations of the marginal posterior of the individual delay $\tau_n$.

We typically used $m = 1,000,000$ and $p = 1$ to generate the figures in the main text and this supplementary material.

### Non-informative hyperpriors for the hierarchical distributed delay model and non-informative priors for its non-hierarchical counterpart

The hierarchical model requires the specification of hyperpriors for all the hyperparameters that describe population variation. In the generative model for the distributed delay case (See Fig. 2b in the main text), we have four pairs of hyperparameters $(a, b)$, one for each of $A$, $B$, $\alpha$, and $\beta$. As information about these hyperparameters may be scarce, especially in real biological systems, specifying non-informative hyperpriors will sometimes be appropriate.

In model implementation, we assumed that the death rates $B_n$ are known and so no longer inferred their population distributions. For the rate parameter, $A_n$, we specified a rational prior with form $\pi(a_A, b_A) = \dfrac{1}{b_A}$. For the delay parameters $\alpha$ and $\beta$, we first considered rational priors of the form $\pi(a_\alpha, b_\alpha) = \dfrac{1}{b_\alpha}$ and $\pi(a_\beta, b_\beta) = \dfrac{1}{b_\beta}$, and afterwards tested changes in estimate accuracy when these are replaced by the MDIP (Eq. S10).

In the comparison done between the hierarchical and non-hierarchical models (See Fig. 3 in main text), we implemented both cases all with non-informative hyperpriors and priors, respectively. For the hierarchical model, we specified a rational joint hyperprior for the pair $(a_A, b_A)$, and MDIP for both the pairs $(a_\alpha, b_\alpha)$ and $(a_\beta, b_\beta)$. Similar to Choi et al. [2020], we specified non-informative gamma priors, $\Gamma(0.001, 0.001)$, for all parameters $A_n$, $\alpha_n$, and $\beta_n$, in the implementation of the non-hierarchical model.

### Supplementary tables and figures

| $\sigma_n^2$ | $(a_\alpha, b_\alpha)$ | $(a_\beta, b_\beta)$ |
|---|---|---|
| 3.5 | $(84, 6)$ | $(10, 5)$ |
| 7 | $(63, 9)$ | $(10, 10)$ |
| 14 | $(35, 10)$ | $(10, 20)$ |

**Table S1**: Hyperparameter values used to generate individual the delay parameters $(\alpha_n, \beta_n)$ that were used to simulate trajectories which served as data for Fig. 2c. In all three cases, the same set of reaction rates $A_n$ and $B_n$ were used, with $A_n \sim \Gamma(8, 0.23)$ and $B_n \sim \Gamma(9, 625)$. In all cases, the mean delay, $\mu_{\tau_n}$, follows a beta prime distribution, $\beta'\left(a_\alpha, a_\beta, 1, \frac{b_\beta}{b_\alpha}\right)$, with mean 7.78 min.

| Hyperparameter | $\sigma_n^2 \approx 3.5$ | | $\sigma_n^2 \approx 7$ | | $\sigma_n^2 \approx 14$ | |
|---|---|---|---|---|---|---|
| $\omega$ | $(\mu_\omega, \sigma_\omega)$ | True value | $(\mu_\omega, \sigma_\omega)$ | True value | $(\mu_\omega, \sigma_\omega)$ | True value |
| $a_\alpha$ | $(81, 3)$ | 84 | $(60, 3)$ | 63 | $(32, 3)$ | 35 |
| $b_\alpha$ | $(6, 3)$ | 6 | $(6, 3)$ | 9 | $(7, 3)$ | 10 |
| $a_\beta$ | $(7, 3)$ | 10 | $(7, 3)$ | 10 | $(7, 3)$ | 10 |
| $b_\beta$ | $(2, 3)$ | 5 | $(7, 3)$ | 10 | $(17, 3)$ | 20 |

**Table S2**: Parameters of the folded normal distribution which served as informative hyperprior for the implementation seen in Fig. S2. In all cases, we used $\rho = 0$.

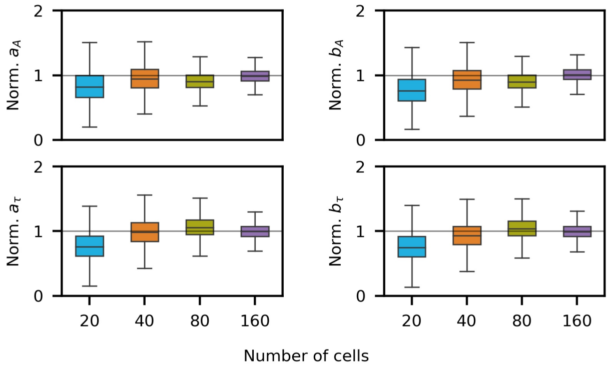


**Fig. S1**: **Increasing the number of cells used in hierarchical inference with fixed delays improved hyperparameter estimates.** We used the hierarchical fixed delay model to infer hyperparameters from simulated trajectories of birth-death processes with fixed birth delays (See 40-minute trajectories in Fig. 1b in main text). Box plots corresponding to hyperparameter posterior distributions obtained using data from an increasing number of cells (from 20 to 160) show the convergence of posteriors to the true hyperparameter values. Estimated values were normalized by dividing with the true hyperparameter values.

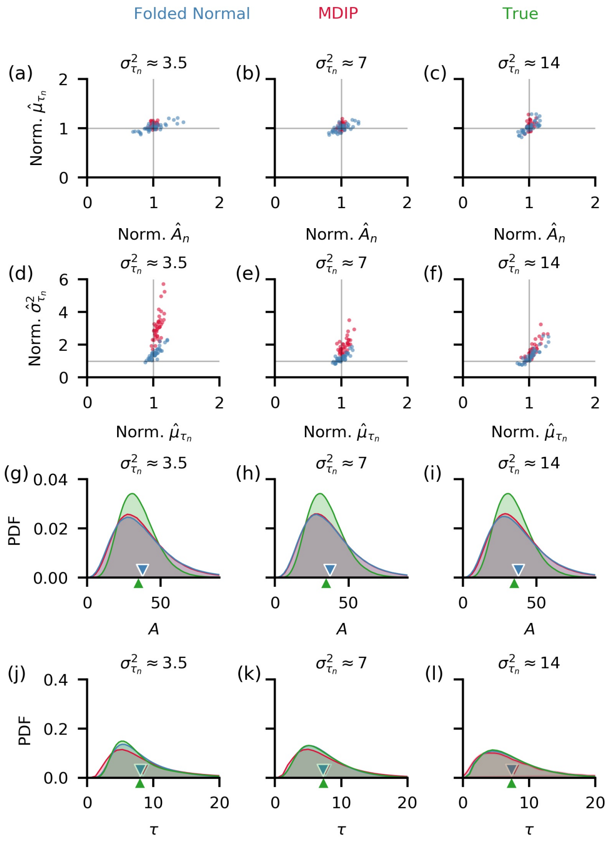


**Fig. S2**: **Informative folded normal delay hyperpriors yielded better estimates of delay parameters which consequently led to better estimates of individual delay variances.** We considered three data sets with different levels of individual delay variability (See Fig. 2 of main text.). We divided the estimates by their true parameter values to facilitate a comparison between different model versions. (a-c) While individual mean delay time estimates, $\hat{\mu}_{\tau_n}$, are similar in both the folded normal and MDIP hyperprior cases, the non-informative MDIP resulted to more accurate production rate, $A_n$, estimates, which may be due to the fact that the strong folded normal hyperpriors were parametrized with values which are smaller when compared to the true generative values (See Table S2.). (d-f) Individual delay variances, however, were significantly better estimated when folded normal delay hyperpriors were specified. (g-l) Population posteriors obtained in both cases closely resemble the true population densities for both the production rate, $A$ (g-i), and delay time, $\tau$ (j-l), across the three data sets considered. The mean of the posteriors (triangular markers) are close to true population means.

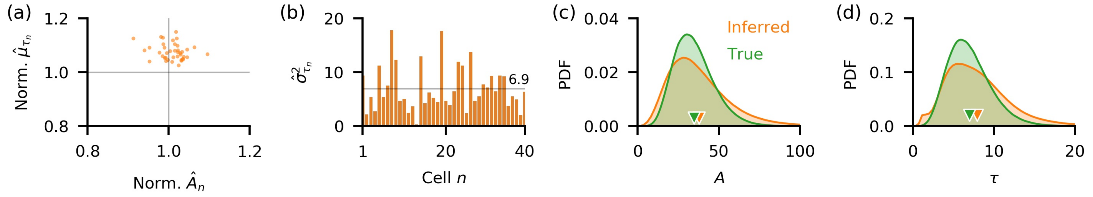


**Fig. S3**: **A distributed delay model with non-informative delay hyperpriors slightly overestimates both the production rate and mean delay times when fit to data with fixed birth delays.** (a) Even with a misspecified generative model, the distributed delay model is able to accurately infer individual parameters of a process with fixed birth delays with a slight overestimation of both the production rates, $A_n$, and mean delay times, $\mu_{\tau_n}$. (b) Since the delay hyperpriors are wide and uninformative, delay variances are largely overestimated with average variance of approximately 6.9 throughout the population, as compared to the true variance 0. (c-d) The slight overestimation of $A_n$ and $\mu_{\tau_n}$ extends to the population distribution whose means (triangular markers) are a bit larger than the true values.

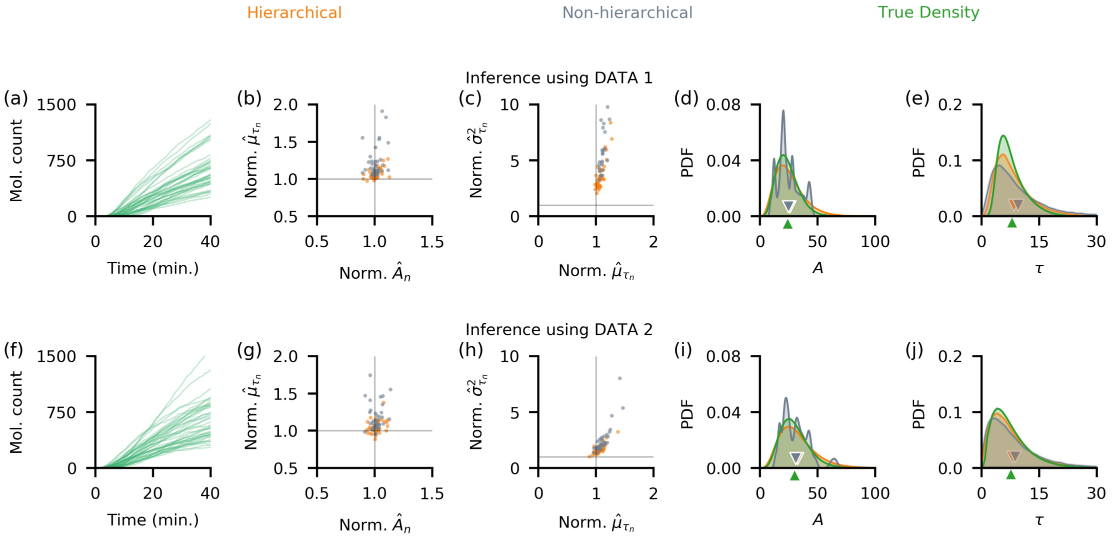


**Fig. S4**: **The hierarchical model consistently outperforms its non-hierarchical counterpart on different parameter and hyperparameter sets.** (a and f) We generated two additional sets of 40 trajectories, each with 40 minutes of observation that is subsampled every minute. The following population distributions were used to generate individual data: $A_n \sim \Gamma(6, 0.25)$, $B_n \sim \Gamma(9, 625)$, $\alpha_n \sim \Gamma(84, 6)$, and $\beta_n \sim \Gamma(10, 5)$ for data 1; and $A_n \sim \Gamma(6, 0.2)$, $B_n \sim \Gamma(9, 300)$, $\alpha_n \sim \Gamma(35, 10)$, and $\beta_n \sim \Gamma(10, 20)$ for data 2. Data 1 has a smaller production rate population mean and narrower individual delay distributions as compared to the data set used in the main text (See Fig. 2c, trajectories with $\sigma^2_{\tau_n} \approx 7$). Data 2, on the other hand, has a larger production rate population mean and wider individual delay distributions. (b and g) While individual production rate estimates, $\hat{A}_n$, are similar in both models, the mean delay times are better estimated with a hierarchical model. (c and h) The same advantage of the hierarchical approach also applies to the estimates of delay variances. (d and i) Although population mean of production rate (triangular markers), $A$, is captured in both approaches, the posterior from the hierarchical model better represent the true density. (e and j) The non-hierarchical model overestimates the population mean of delay times (triangular markers) while the hierarchical model gives a more accurate estimate.

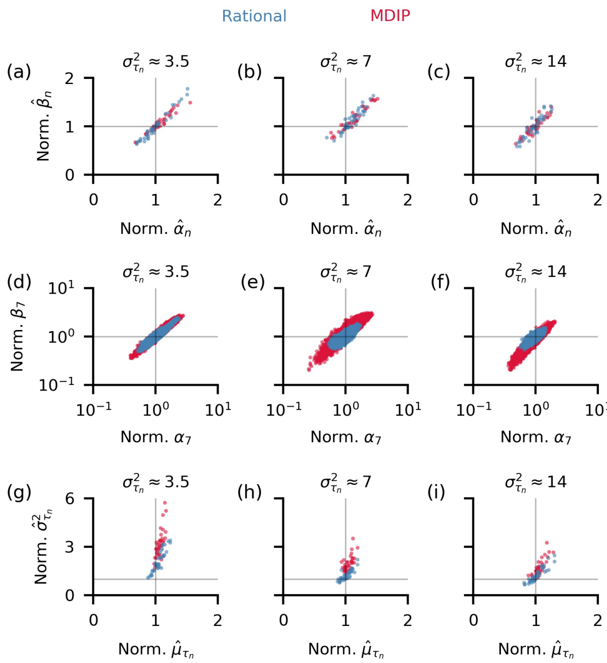


**Fig. S5**: **Variance of individual delay distributions are better captured using rational delay hyperpriors but this advantage disappears as true delay distributions become wider.** In the distributed delay model, we used two different non-informative delay hyperparameter distributions in three different implementations: rational priors and the MDIP. (a-c) Estimates of individual delay parameters $(\alpha_n, \beta_n)$ are similar in both choices of non-informative priors. (d-f) Samples of individual estimates of the delay parameters $\alpha$ and $\beta$, for cell 7. The posterior distribution over the parameter shows a strong correlation between the two. Similar correlations are observed for all cells, both when using the hierarchical, and non-hierarchical model. (g-i) Errors in the estimates of $(\alpha_n, \beta_n)$ lead to the overestimation of delay variances in model implementations using the rational and MDIP delay hyperpriors. While the errors in the estimates remained small in the case of the rational hyperpriors in all three data sets considered, the estimates improved for the MDIP case, as true individual delay variances become larger.

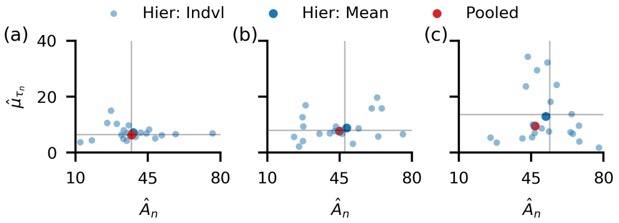


**Fig. S6**: **While pooling of data is good for estimating mean parameter values for data with little variation across the population, errors may increase as cells become more different.** Twenty trajectories accounting for 20-minute observations of a delayed stochastic birth-death process served a data in this comparison. In order of increasing variability, both in terms of mean delays and production rates, across the population, data 1 (a) has the least variability, next is data 2 (b), while data 3 (c) has the largest. The following population distributions were used to generate individual data: $A_n \sim \Gamma(8, 0.23)$, $B_n \sim \Gamma(9, 625)$, $\alpha_n \sim \Gamma(63, 9)$, and $\beta_n \sim \Gamma(10, 10)$ for data 1; $A_n \sim \Gamma(8, 0.16)$, $B_n \sim \Gamma(9, 625)$, $\alpha_n \sim \Gamma(7, 1)$, and $\beta_n \sim \Gamma(5, 5)$ for data 2; and $A_n \sim \Gamma(8, 0.16)$, $B_n \sim \Gamma(9, 625)$, $\alpha_n \sim \Gamma(3.3, 0.6)$, and $\beta_n \sim \Gamma(2, 2.5)$ for data 3. As the variability increases, the estimates from model with data pooling migrate farther away from the true population means (vertical and horizontal lines in each plot), while the means of the hierarchical model estimates remain accurate.

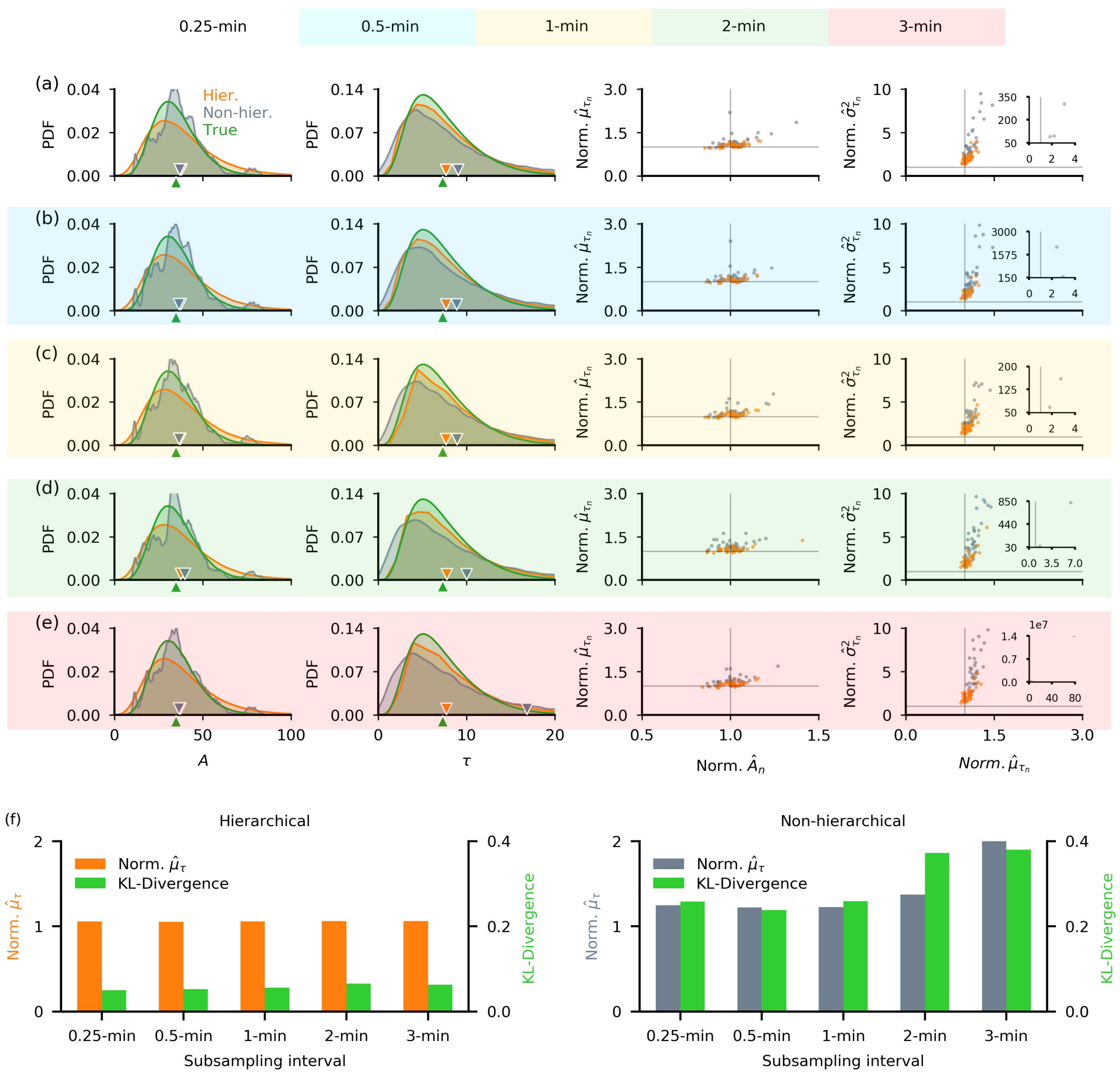


**Fig. S7**: **A non-hierarchical model is more sensitive to changes in sampling frequency than a hierarchical model.** We implemented the hierarchical model and its non-hierarchical counterpart using 20 minutes of subsampled data (see $\sigma_n^2 \approx 7$ trajectories in Fig. 2c in the main text) with decreasing sampling frequency (from 4 per min, i.e. 0.25-min subsampled, to 1/3 per min, i.e. 3-min subsampled). Although population means of the production rate, $A$, are very similar in both models across all subsampling schemes (triangular markers in a-e 1st column), the accuracy of the estimate of the delay distribution mean from the non-hierarchical model (grey) decreased with sampling frequency while those from the hierarchical model (orange) exhibited a similar level of accuracy (triangular markers in a-e 2nd column). Across all data subsets we considered, the hierarchical model individual parameter estimates for $A_n$, $\mu_{\tau_n}$ (a-e 3rd column), and $\sigma^2_{\tau_n}$ (a-e 4th column) exhibited small deviations. Estimates from the non-hierarchical model, on the other hand, reduced in accuracy especially in terms of $\sigma^2_{\tau_n}$ (a-e 4th column) when we decreased the sampling frequency, with extreme outlying estimates produced at low sampling frequencies (a-e 4th column inset). (f) A comparison of population delay distributions showed that the hierarchical model produced a mean delay estimate (left - orange bars ) that is consistently accurate, together with a population posterior with low KL-divergence from the posterior to the true density (left - green bars) that barely changed between the data subsets we considered. Decrease in sampling frequency resulted in reduced accuracy of the population mean delay estimate from the non-hierarchical model (right - grey bars). The non-hierarchical delay posterior also exhibited KL-divergence that increased with the subsampling interval (right - green bars).

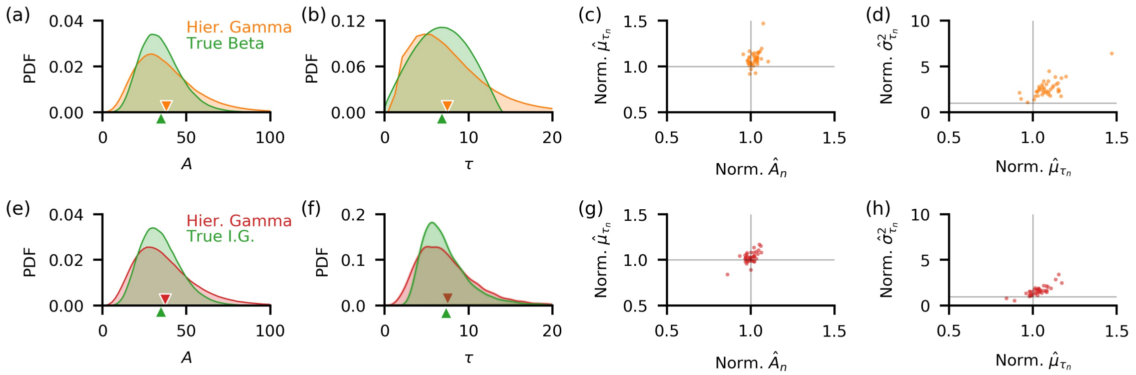


**Fig. S8**: **The hierarchical model provides accurate estimates even when the delay distribution is mismatched.** We fit the hierarchical model with gamma distributed individual cell birth delays to data generated using beta (a-d) and inverse-gamma (e-h) distributions for the same. The gamma distribution specified in our model has infinite support and decays exponentially, while beta distribution has compact support and the inverse-gamma distribution is heavy-tailed. Even when the delay distributions in the model and data are not matched, population posteriors obtained in both cases closely resemble the true population densities for both the production rate, $A$ (a and e), and delay time $\tau$ (b and f). The mean of the posteriors (triangular markers) are close to true population means. Individual estimates of the mean delay, $\mu_{\tau_n}$ (c and g), are accurate, while delay variances, $\sigma^2_{\tau_n}$ (d and h), are slightly overestimated, as in when the distributions in the model and data are matched (see Fig. 3b in main text).

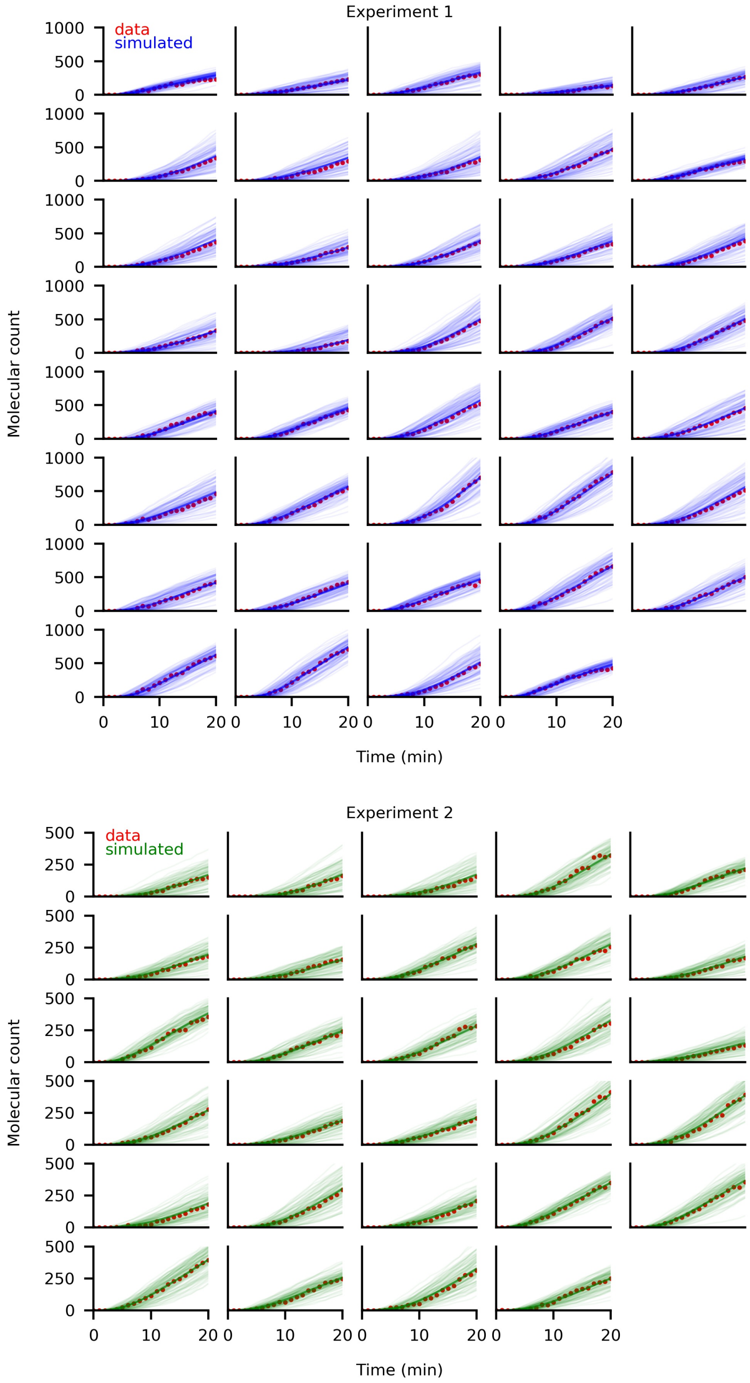


**Fig. S9**: **Simulated realizations with estimated parameters fit individual YFP trajectories.** We simulated 100 trajectories for each cell by sampling the parameters from the 95% high density interval (HDI) of the posterior distributions, using the delayed Gillespie algorithm [Barrio *et al.*, 2006]. The mean of the realizations (solid lines), per cell, fit the experimental data very well.

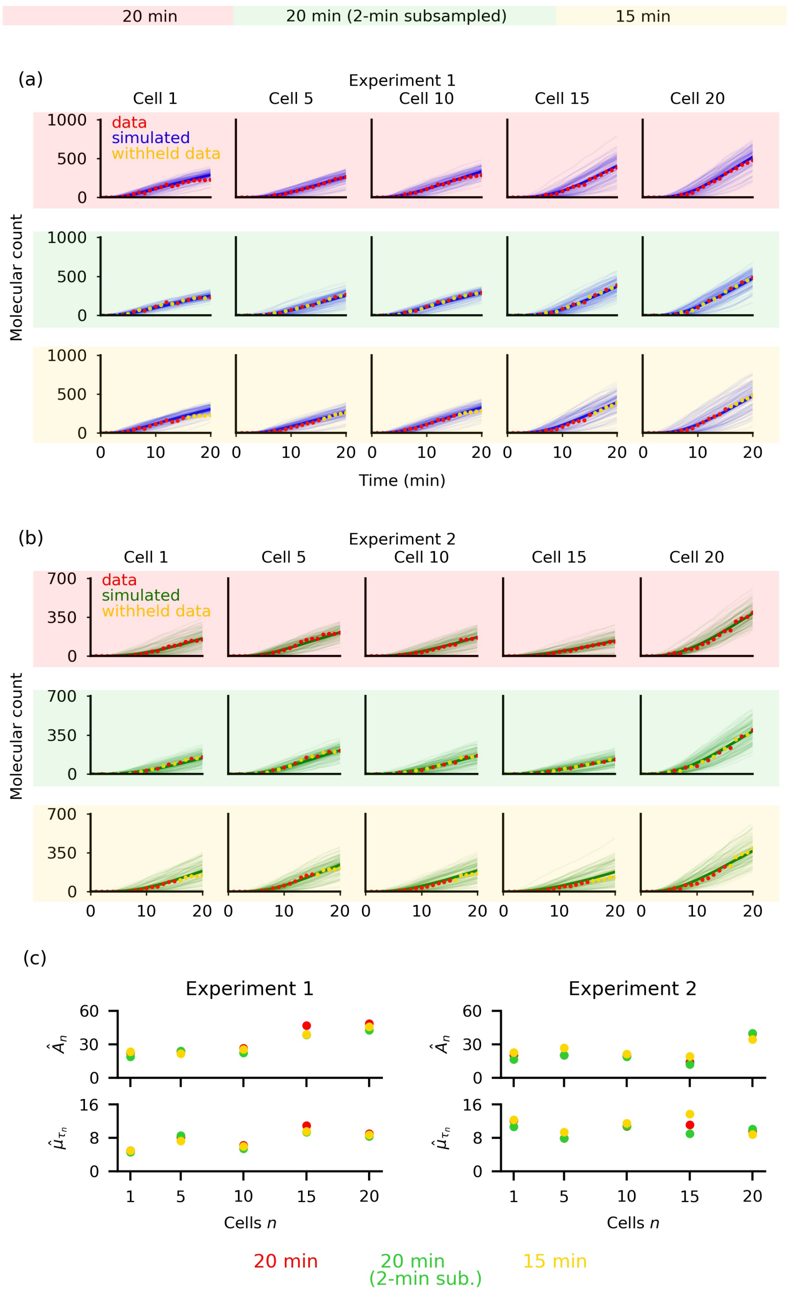


**Fig. S10**: **Simulated realizations with estimated parameters from the hierarchical distributed delay model fit individual YFP trajectories even when some data points were withheld during inference.** Setting the death rates, $B_n$, to their true values during inference, we fit the model to subsets of the experimental data: full 20 min (red background), 20 min with data subsampled at 2-min intervals (green background), and the first 15 min (yellow background) data. Simulated trajectories for experiments 1 (a) and 2 (b) using the inferred parameters across in all settings we considered fit data well. (c) Inference results of five randomly selected cells are shown. Individual estimates of production rates, $A_n$, and mean delay time, $\mu_{\tau_n}$, showed small deviations with changes in the data set indicating that inference is robust. See Fig. **S13** for the analysis of the inference using all cells.

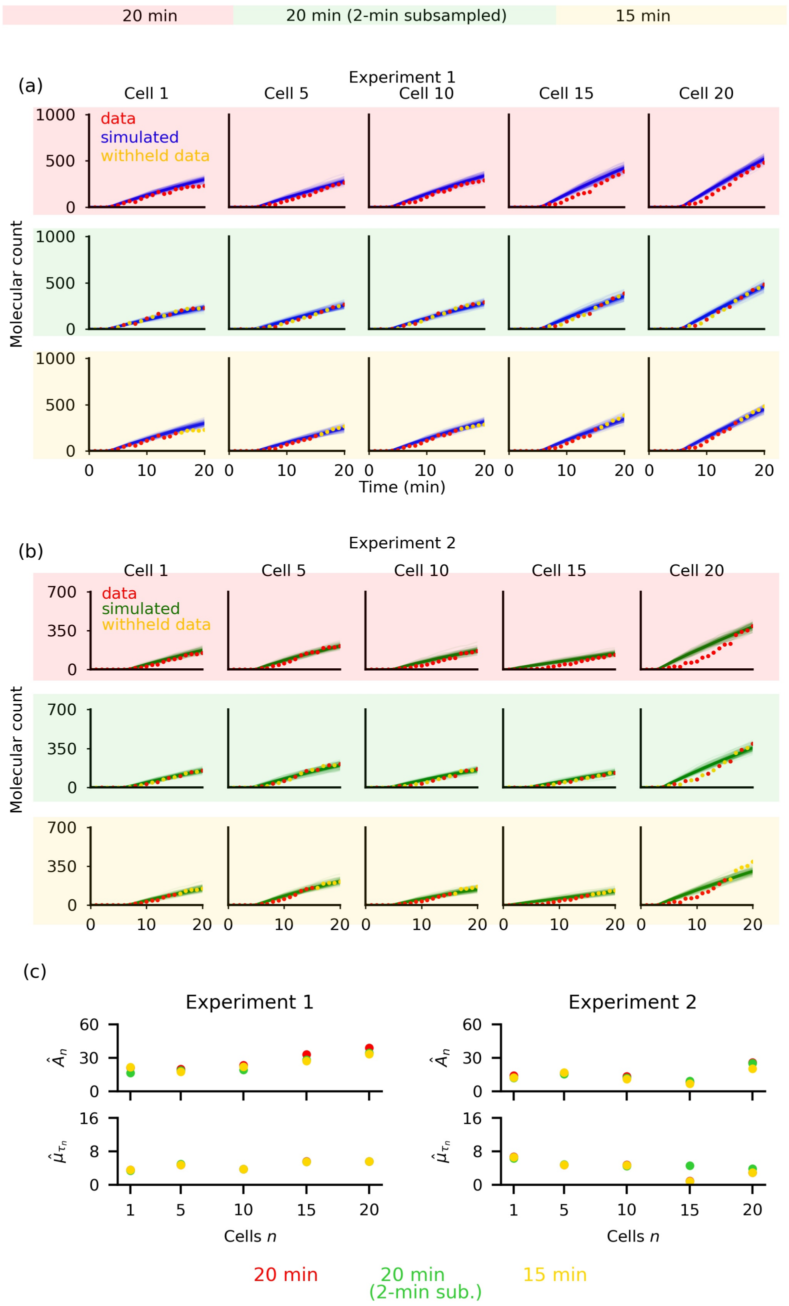


**Fig. S11**: **Simulated realizations with estimated parameters from the hierarchical *fixed* delay model do not exhibit the sigmoidal trajectories that characterize the YFP data.** Setting the death rates, $B_n$, to their true values during inference, we fit the fixed delay model to subsets of the experimental data: full 20 min (red background), 20 min with data subsampled at 2-min intervals (green background), and the first 15 min (yellow background) data. In both experiments 1 (a) and 2 (b), simulated trajectories closely matched the initial and final data points but deviated from the data in the middle of the trajectory. (c) Inference results of five randomly selected cells are shown. Individual estimates of production rates, $A_n$, and mean delay time, $\mu_{\tau_n}$, showed small deviations with the change in data set. See Fig. **S13** for the analysis of the inference using all cells.

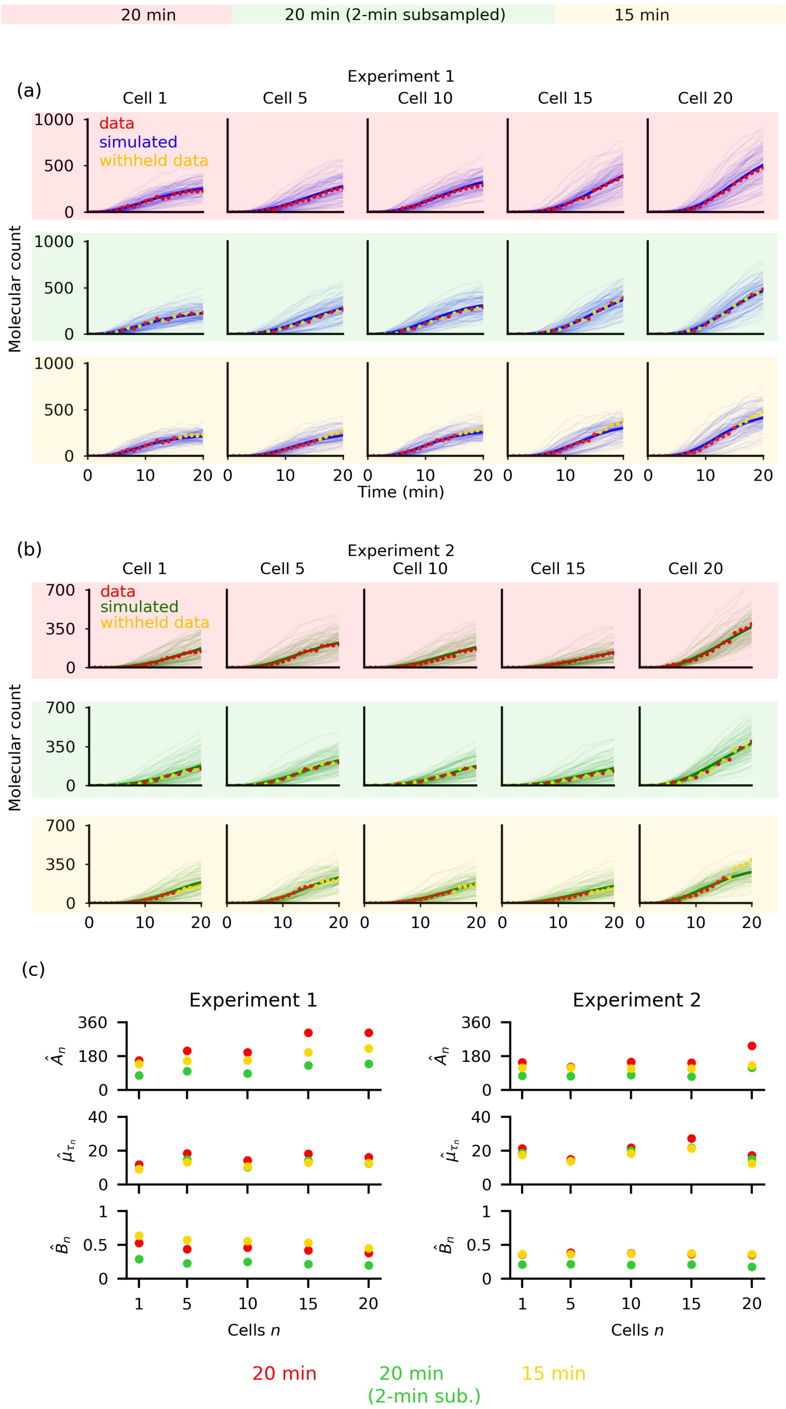


**Fig. S12**: **Full parameter set estimation using the hierarchical distributed delay model resulted in unrealistically large estimates that produced simulated realizations which fit individual YFP trajectories well.** We fit the model to subsets of the experimental data: full 20 min (red background), 20 min with data subsampled at 2-min intervals (green background), and the first 15 min (yellow background) data. Simulated trajectories for experiments 1 (a) and 2 (b) using the inferred parameters across all settings we considered fit data well. (c) Inference results of five randomly selected cells are shown. Individual estimates of production rates, $A_n$, mean delay time, $\mu_{\tau_n}$, and death rate, $B_n$, all are unrealistically large. See Fig. **S13** for the analysis of the inference using all cells.

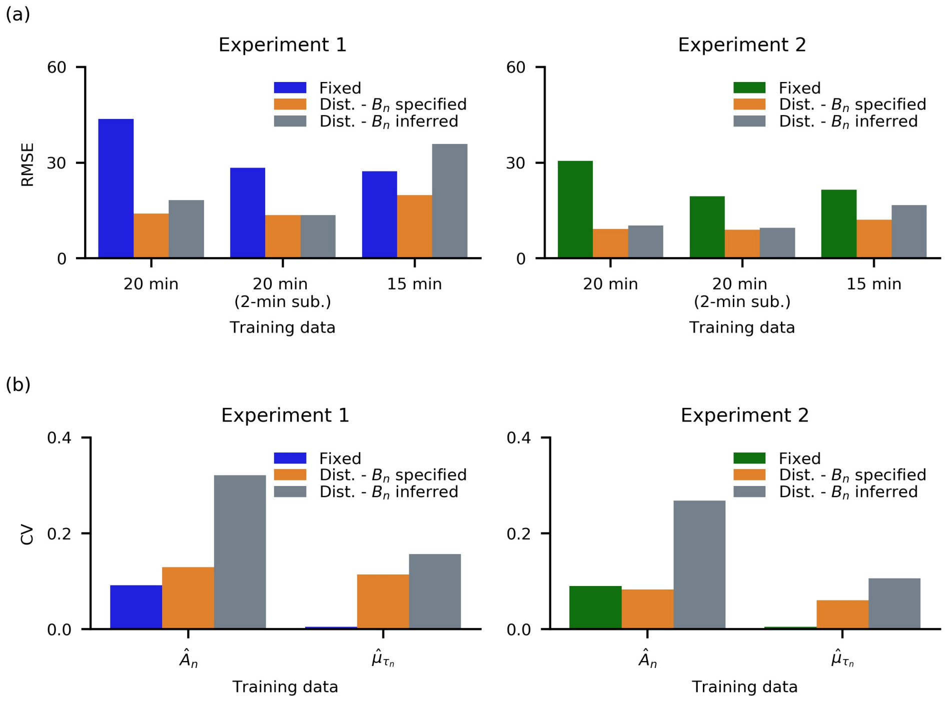


**Fig. S13**: **Fixed delay and unspecified death rate lead to underfitting and overfitting respectively.** (a) We computed the root mean square error (RMSE) of the mean simulated trajectories (see Fig. S9) from the experimental data per individual cell, and averaged over all cells. In both experiments 1 (left) and 2 (right), the RMSE remained low with small changes in the case of the distributed delay model where $B_n$ was specified. In the case of the fixed delay model the error unexpectedly increased with the amount of data used to infer the parameters and hyperparameters, indicating a larger bias. Inference of the full parameter set (including $B_n$) using the distributed delay hierarchical model resulted in larger RMSEs compared to when $B_n$ was specified. (b) We computed the coefficient of variation (CV) of the parameter estimates (see Fig **S10**c, **S11**c, **S12**c ) across the different data subsets per individual, then averaged over all individuals. The fixed delay model showed the least variation among the models then followed by the distributed model with $B_n$ specified. The distributed model where all parameters were inferred exhibited the largest variation in all parameters.
